# Effects of Data Transformation and Model Selection on Feature Importance in Microbiome Classification Data

**DOI:** 10.1101/2023.09.19.558406

**Authors:** Zuzanna Karwowska, Oliver Aasmets, Estonian Biobank research team, Tomasz Kosciolek, Elin Org

## Abstract

Accurate classification of host phenotypes from microbiome data is essential for future therapies in microbiome-based medicine and machine learning approaches have proved to be an effective solution for the task. The complex nature of the gut microbiome, data sparsity, compositionality and population-specificity however remain challenging, which highlights the critical need for standardized methodologies to improve the accuracy and reproducibility of the results. Microbiome data transformations can alleviate some of the aforementioned challenges, but their usage in machine learning tasks has largely been unexplored. Our aim was to assess the impact of various data transformations on the accuracy, generalizability and feature selection by analysis using more than 8,500 samples from 24 shotgun metagenomic datasets. Our findings demonstrate the feasibility of distinguishing between healthy and diseased individuals using microbiome data with minimal dependence on the algorithm and transformation selection. Remarkably, presence-absence transformation performed comparably well to abundance-based transformations, and only a small subset of predictors is crucial for accurate classification. However, while different transformations resulted in comparable classification performance, the most important features varied significantly, which highlight the need to reevaluate machine-learning based biomarker detection. Our research provides valuable guidance for applying machine learning on microbiome data, offering novel insights and highlighting important areas for future research.

## Introduction

Human microbiome carries a vast amount of information that can be used to improve the understanding of our functioning and be potentially used to improve clinical practice and public health. Harnessing this information on the other hand is not trivial due to the complexity of the microbial ecosystem, which comprises hundreds of species and involves intricate interactions between the ecosystem members. Similarly to other fields of biology, machine learning (ML) approaches have become pivotal in microbiome research as they can inherently account for the high dimensionality and versatile data types. Predicting an outcome based on the taxonomic or functional profile is perhaps the most widespread use of ML in the microbiome field, however thanks to its versatility, ML is used for taxonomic assignment, functional profiling and others (Marcos-Zambrano et al. 2021). ML has been successfully used to build classification models for diseases such as colorectal cancer (Wirbel et al. 2019) and pancreatic cancer (Kartal et al. 2022), but also for predicting the disease outcome in the future such as for liver diseases (Liu et al. 2022), type 2 diabetes (Ruuskanen et al. 2022) or all cause mortality (Salosensaari et al. 2021).

Currently, analysis of the microbiome data lacks standards and best approaches are yet to be identified (Kubinski et al. 2022; Hernández Medina et al. 2022). For example, differential abundance analysis, a common analysis step to identify members of the microbiome whose abundance is different between the study groups, has been shown to produce remarkably varying results depending on the analysis methodology used (Nearing et al. 2022). Such conflicting results can be explained by the unique properties of microbiome data, such as compositionality, high dimensionality, and high sparsity, which pose challenges for standard statistical methods and by the observation that many DA methods evaluate tests on very different estimates (Gloor et al. 2017). To address these limitations, various data transformations like total-sum-scaling (TSS), arcsine-square-root (aSIN), and log-ratio transformations such as centered-log-ratio (CLR), isometric-log-ratio (ILR) or additive log-ratio (ALR) are commonly employed in microbiome research (Ibrahimi et al. 2023). However, the impact of data transformations on prediction and classification tasks employing machine learning algorithms remains poorly understood.

Recently, Giliberti *et al*. carried out an extensive analysis to compare the performance of models based on the presence-absence of microbes and TSS scaling (Giliberti et al. 2022). Intriguingly, they found that presence-absence of the microbes as features in a predictive model leads to equivalent predictive performance. However, there are indications that other data transformations, especially log-ratio based transformations can outperform the TSS in predictive tasks. For example, CLR has been shown to improve the prediction accuracy over TSS (Kubinski et al. 2022; Delgado et al. 2019). Nevertheless, in light of the results by Giliberti *et al*., it remains unclear whether the aforementioned data transformations can improve the prediction accuracy over presence-absence.

Here, we systematically evaluate the impact of various data transformations on the binary classification performance (e.g. distinguishing healthy and diseased individuals) to determine the optimal modeling strategies for shotgun metagenomics data. We employ eight data transformations in combination with threeML algorithms (random forest, extreme gradient boosting and elastic net) and assess their performance on 24 metagenomic datasets across various disease outcomes to ensure an unbiased and robust assessment. In addition, we investigate how the selection of the data transformation impacts the external generalizability and feature selection, which is essential for biomarker discovery.

## Results

### Study design

To investigate the impact of the data transformations on the binary classification performance, we used publicly available shotgun metagenomic sequencing datasets present in the *curatedMetagenomicData* R package (version 3.6.2), which encompass more than 6,000 samples across different populations and phenotypes (Pasolli et al. 2017). In our analysis, we focused on stool metagenomic datasets with a primary phenotype available and that had at least 50 cases and 50 controls (**Supplementary Table 1**, **Methods**). Additionally, we used the metagenomic data from the Estonian Microbiome Cohort (EstMB), which is coupled with rich phenotype data (N = 2,509) (Aasmets et al. 2022). **Figure 1a** shows the study design and study objectives. Firstly, each metagenomic dataset was transformed using eight data transformations, which are commonly applied in the microbiome field. The transformations included presence-absence transformation (PA), relative abundance transformation (total sum scaling, TSS), logarithm of TSS, arcsine square root transformation (aSIN), and four compositional transformations (centered log-ratio (CLR); robust centered log-ratio (rCLR) isometric log-ratio (ILR) and additive log-ratio (ALR)). For sensitivity analysis, we additionally rarefied the datasets before applying data transformations. The transformed datasets were then used in a binary classification setting using three learning algorithms, random forest (RF), extreme gradient boosting (XGB)and elastic net (ENET). Our primary objective was to assess the classifier performance across the data transformations within the analyzed datasets (within-study (WS) setting). Secondly, we aimed to assess the impact of data transformations in different analytical scenarios. In addition to the within-study setting, we evaluated the external generalizability of the models by carrying out a leave-one-study-out cross-validation for colorectal cancer (CRC; 11 datasets, **Supplementary Table 1**) and obesity (subjects with BMI > 30; BMI30; 5 datasets, **Supplementary Table 1**). Furthermore, we analyzed whether the data transformations benefit from a larger sample size and whether the corresponding model performance is dependent on the number of predictors by altering the data dimensionality and evaluating the effects on the classification performance. Lastly, we analyzed the features selected by the models to gain additional insights about how data transformations impact the conclusions of the ML analysis.

**Figure 1.**
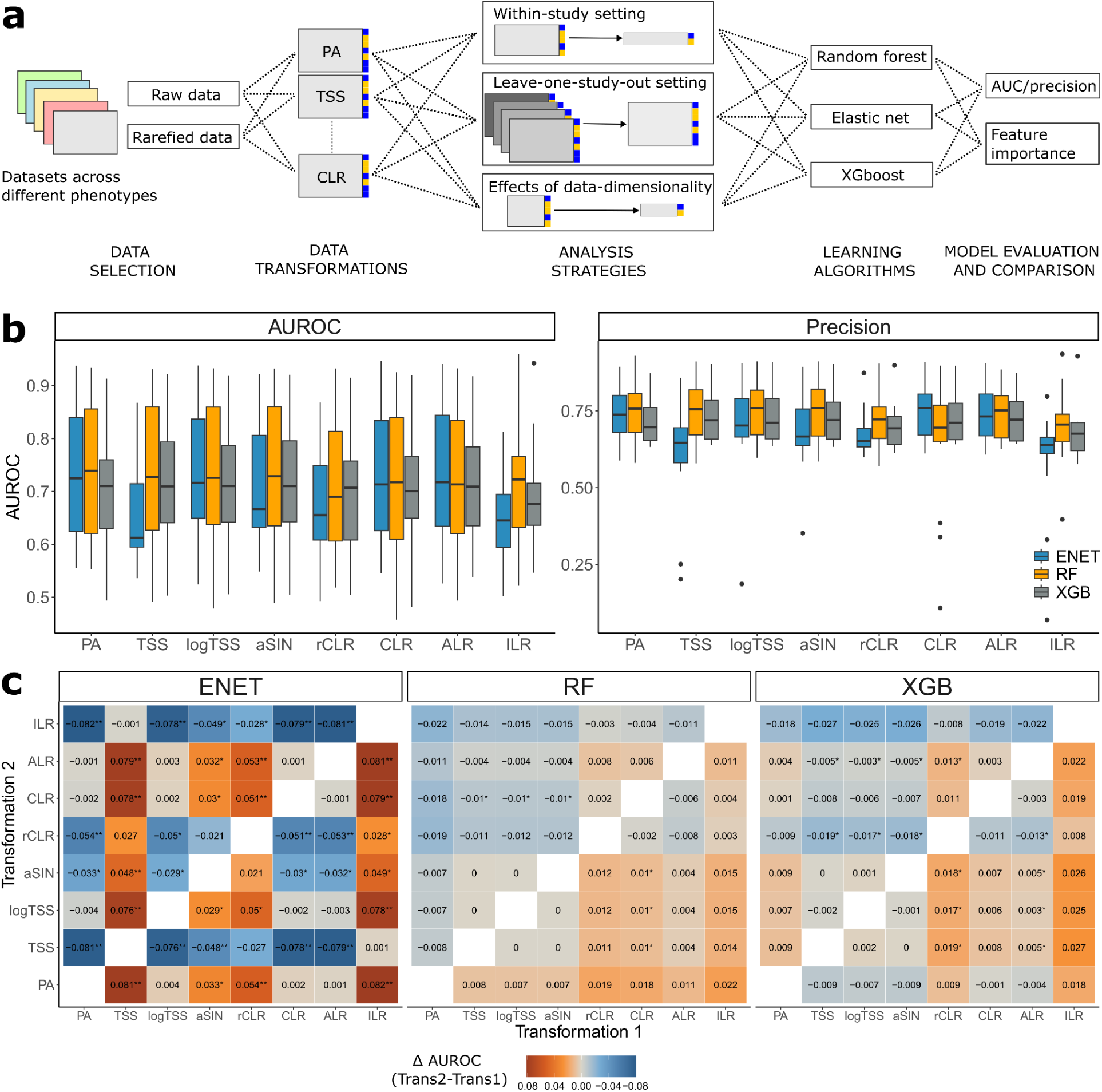
STUDY DESIGN AND CLASSIFICATION RESULTS. **a** - Overview of the study design. **b** - Classification performance (AUROC and precision) in the within-study setting for every data transformation and algorithm. **c -** Statistical analysis results between the used data transformations (Wilcoxon signed-rank test) for elastic net (ENET),random forest (RF) and extreme gradient boosting (XGB); Values and colors correspond to the differences in AUROC between Transformation 2 and Transformation 1; * indicates a nominally statistically significant difference in AUROC (Wilcoxon signed-rank test, p-value ≤ 0.05), ** indicates a statistically significant difference in AUROC after correction (FDR ≤ 0.05). Abbreviations: ENET - elastic net logistic regression; RF - random forest; XGB - extreme gradient boosting, XGBoost; PA - presence-absence; TSS - total-sum scaling; logTSS - logarithm of TSS; aSIN - arcsine square root; CLR - centered log-ratio; rCLR - robust CLR; ALR - additive log-ratio; ILR - isometric log-ratio

### Overview of classifier performance dependence on data transformations

Our primary aim was to analyze whether the machine learning performance in a binary classification task depends on the selected data transformation and whether the data transformations are leveraged differently by distinct ML algorithms. The comparison of the area under the receiver operating characteristic (AUROC) for random forest (RF), extreme gradient boosting (XGB) and elastic net logistic logistic regression (ENET) by data transformations are shown in **Figure 1b-c**. On average, the performance of ENET was significantly lower than RF and XGB when TSS was used as a data transformation (Wilcoxon signed-rank test, FDR ≤ 0.05). Similarly, RF outperformed ENET with ILR and rCLR and RF outperformed XGB with PA. With other data transformations, differences between the algorithm performances were not statistically significant (**Figure 1b**). For ENET, XGB and RF, rCLR performed significantly worse than several other data transformations indicating that rCLR is not fit for ML purposes. Similarly, ILR transformation led to significantly lower performances compared to other data transformations, which, however, was not statistically significant for RF and XGB. Other than that, we did not identify significant differences in classification performances between the data transformations that were universal for thelearning algorithms and across different datasets (**Figure 1b-c, Supplementary Figure 1**). For ENET, ILR, rCLR, aSIN and TSS resulted in inferior performance compared to the other data transformations (Wilcoxon signed-rank test, FDR ≤ 0.05). Importantly, PA for ENET was better or equivalent to other data transformations in terms of predictive performance. In contrast, RF and XGB did not exhibit as pronounced differences in AUROC between different data transformations, although the usage of PA with RF led to better classification performance for RF when compared to ILR, CLR, rCLR and ALR (**Figure 1b-c**). Similarly, RF in combination with TSS, logTSS and aSIN outperformed CLR (nominal significance, p-value ≤ 0.05). For XGB, ALR, aSIN, TSS and logTSS led to better performance than rCLR, other differences were not statistically significant (nominal significance, p-value ≤ 0.05). As a sensitivity analysis, we carried out rarefaction before applying the data transformations. In this scenario, we observed highly similar results to the non-rarefied case with PA leading to optimal predictive performance (**Supplementary Figures 2,3**). On average, the performance of the rarefied data was lower compared to the unrarefied data for aSIN (FDR = 0.0062), TSS (FDR = 0.0083), logTSS(FDR = 0.0012) and ALR (FDR = 0.0155) inidcating that for binary classification on the shotgun metagenomics data, rarefaction is not necessary. Thus, our results are consistent with the results by Giliberti *et al (Giliberti et al. 2022)* showing that presence-absence (PA) leads to equivalent or even better classification performance as compared to using relative abundances. Moreover, our results show that the same can be concluded for other commonly used data transformations.

### Data transformation effects in different analytical scenarios

We were surprised that no significant improvement in classification performance was observed when abundance-based transformations were used instead of PA. To understand whether the data transformations could give advantage in other analytical scenarios, we conducted several follow-up analyses. Firstly, we assessed how the sample size and number of features in the initial dataset influence the classification performance. We hypothesized that some data transformations may lead to better performance in certain sample size/data dimensionality settings. For that, we applied different prevalence thresholds to the microbial taxa on the publicly available metagenomics datasets and on the Estonian Microbiome Cohort (EstMB) dataset (N = 2,509) before carrying out the classification task. For the EstMB dataset, we additionally subsampled the cases and controls of obesity (BMI > 30) and antibiotics usage (90 days from sample collection) (20%, 40%, 60% and 80% of the initial number of cases and controls) to study the impact of varying sample size and number of predictors at once. As expected, we observed that larger sample sizes and the inclusion of less prevalent taxa leads to improved classification performance (**Figure 2a**, **Supplementary Figure 2**). Nevertheless, we found no substantial interactions on the classification performance between the data transformations, sample size and the number of features.

**Figure 2.**
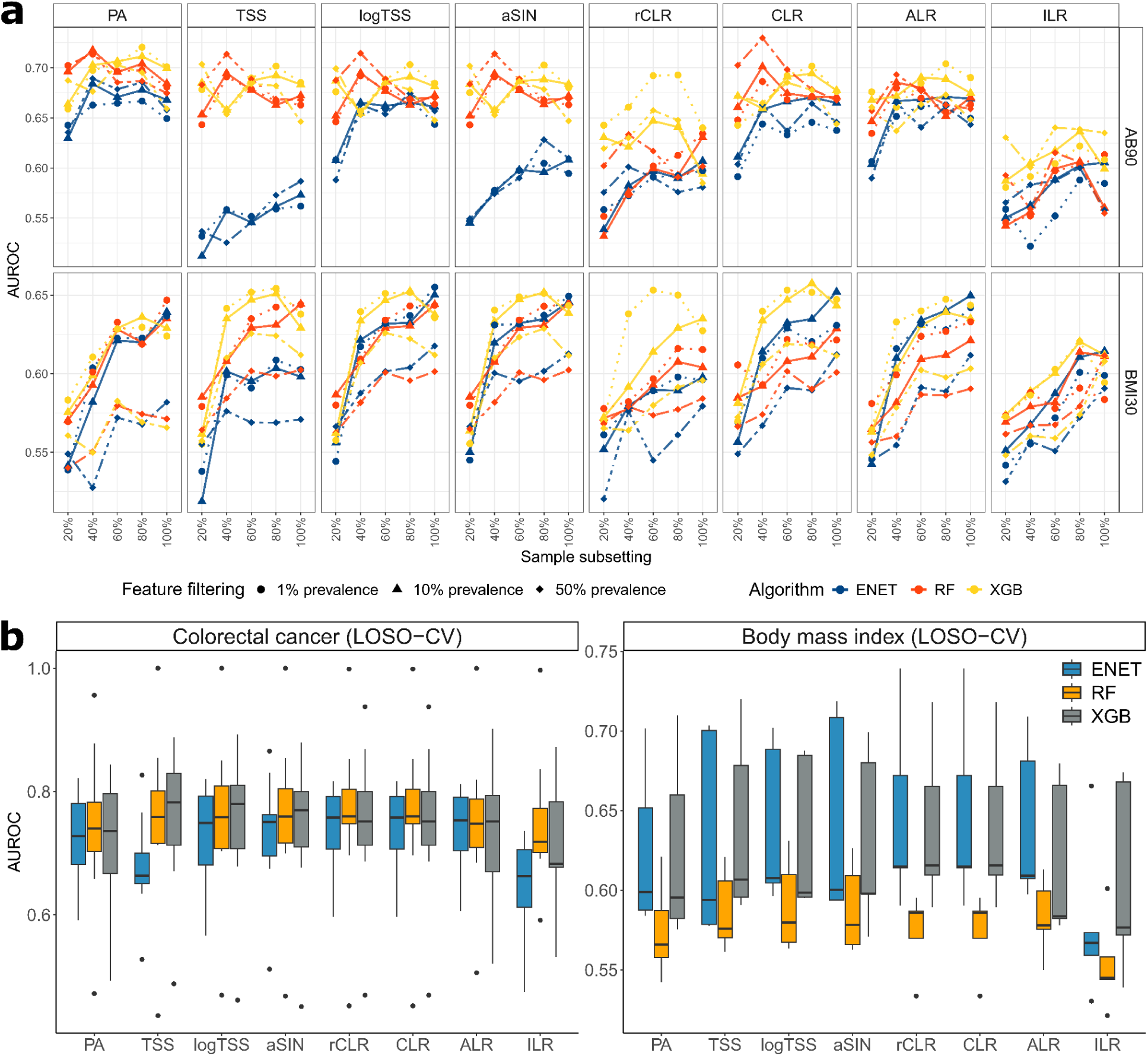
IMPACT OF DATA TRANSFORMATIONS ON CLASSIFICATION PERFORMANCE UNDER DIFFERENT SCENARIOS. **a** - Classification performance across varied data dimensions and sample sizes for random forest (RF), extreme gradient boosting, XGBoost (XGB) and elastic net logistic regression (ENET) and every transformation. **b** - Transformation and Model Outcomes in a Leave-One-Study-Out cross-validation (LOSO-CV) setting. Abbreviations: ENET - elastic net logistic regression; RF - random forest; XGB - extreme gradient boosting, XGBoost; PA - presence-absence; TSS - total-sum scaling; logTSS - logarithm of TSS; aSIN - arcsine square root; CLR - centered log-ratio; rCLR - robust CLR; ALR - additive log-ratio; ILR - isometric log-ratio; AB90 - antibiotics use within 90 days from sample collection; BMI30 - obesity as defined by BMI > 30.

Next, we assessed whether data transformations impacted the model’s ability to generalize to unseen data by measuring its classification performance on an external dataset. To do this, we employed a leave-one-study-out (LOSO) validation method for both colorectal cancer (CRC) and obesity defined by BMI > 30 (BMI30) datasets. This involved training a model on a combined set of samples from all studies except one and evaluating its performance on the omitted study. Similarly to the within-sample setting, we observed no significant improvement in the model generalizability when employing abundance-based data transformations (**Figure 2b**). Thus, our analysis indicates that in terms of classification performance, presence-absence is usually a good option and should be considered as an alternative to the abundance-based transformations.

### Feature importance

As no data transformation could consistently be considered superior in terms of classification performance and several data transformations led to equivalent performance, we were interested in how different data transformations impact feature selection and feature importance. To assess feature importance, we calculated mean absolute SHapley Additive exPlanations (SHAP) values for each dataset and for each microbe. SHAP values are a method used in machine learning to explain the contribution of each feature to the prediction of a model (Lundberg and Lee 2017). Focusing on predictors with a non-zero mean absolute SHAP value, we found that the number of selected predictors was highly transformation-specific. Compositional transformations ALR,CLR and rCLR selected more predictors, particularly when used with RF (**Figure 3a**). However, regardless of the transformation, only a small subset of features (∼ below 25 features) held significant importance (**Supplementary Figure 5**). To validate this, we built classifiers for obesity, depression and antibiotics use on the PA-transformed EstMB cohort data that cumulatively use only the most significant features. Surprisingly, just 10 most significant microbial predictors for antibiotics, 25 for depression, and 75 for BMI resulted in comparable classification performance when compared to models using the full microbiome profile, with performance decreasing as more features were added (**Figure 3b, Methods**). We believe the decline in model performance with additional features is due to the unique characteristics of gut microbiome data. Its compositional, high-dimensional nature can cause overfitting, especially with small sample sizes. Using fewer key features helps reduce noise and improves model accuracy. Dimensionality reduction methods such as principal component analysis (PCA), non-metric multidimensional scaling (NMDS), or non-negative matrix factorization (NMF) can address these challenges by transforming highly correlated features into orthogonal vectors. However, these techniques come with limitations, including the need for careful data transformation, appropriate pseudocount use, and accounting for phylogenetic interactions. Additionally, understanding which features drive classification becomes harder to interpret with these methods..(Armstrong et al. 2022)

**Figure 3.**
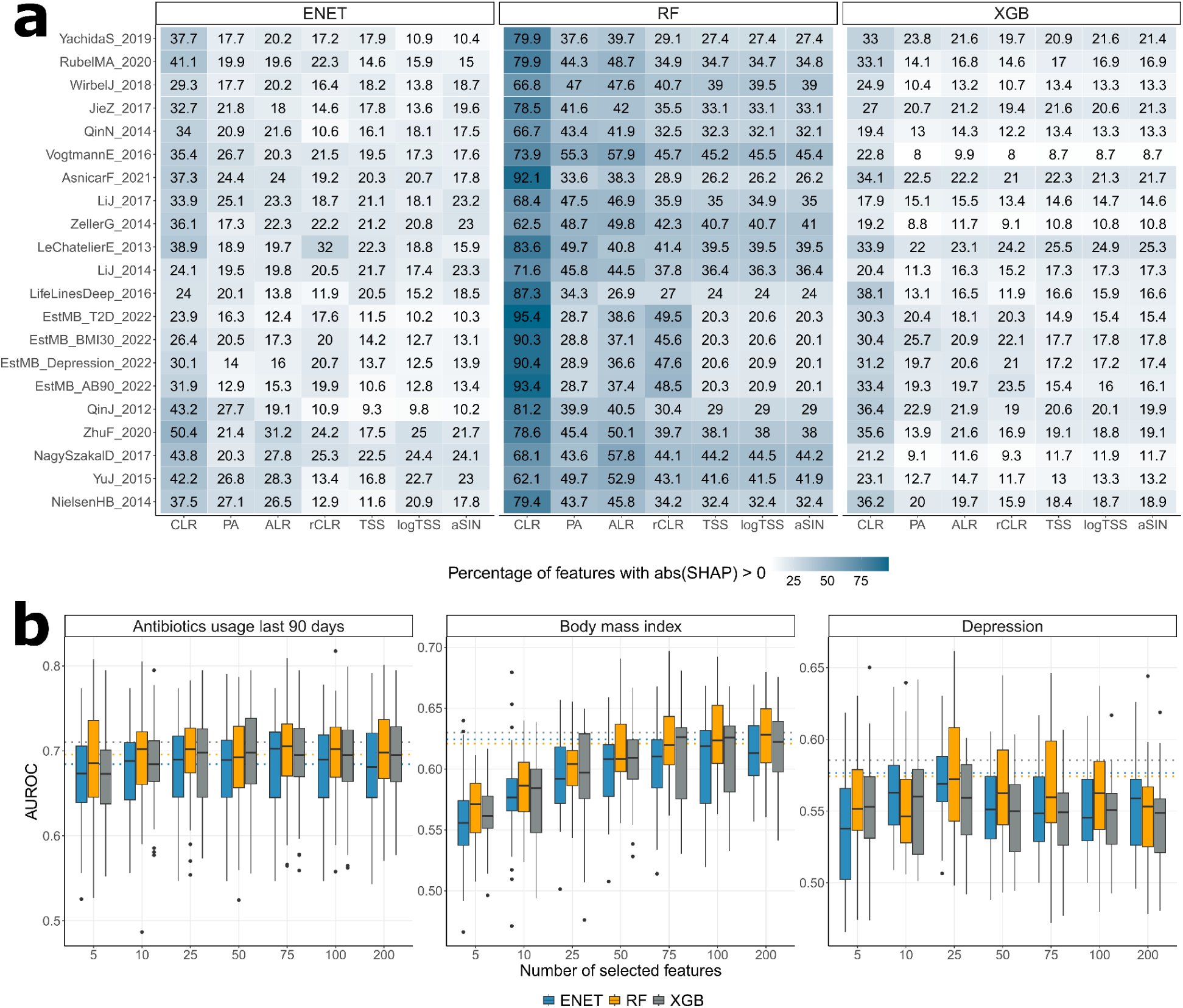
FEATURE SELECTION AMONG DATA TRANSFORMATIONS. **a** - Number of Selected Features: Proportion for ENET, XGB and RF. **b** - XGB, ENET and RF Classification Performance on presence-absence (PA) data using top-N features. The dashed line indicates classifier performance on all features.

Next, we examined the features with the highest predictive value across the transformation-algorithm combinations. Building on our earlier results highlighting a small subset of features with high SHAP values, we first focused on the overlap among the top 25 predictors exhibiting the highest mean absolute SHAP values (**Figure 4a**). For ENET, TSS, rCLR and to lesser extent aSIN exhibited lower overlap with other transformations, while the highest agreement was found between PA, CLR, ALR and logTSS. The overlap between the top predictors for RF was remarkably higher with TSS, aSIN and logTSS showing almost perfect correspondence. The top predictors for XGB closely aligned with those from RF, showing strong similarity across TSS, aSIN, and logTSS transformations.

**Figure 4.**
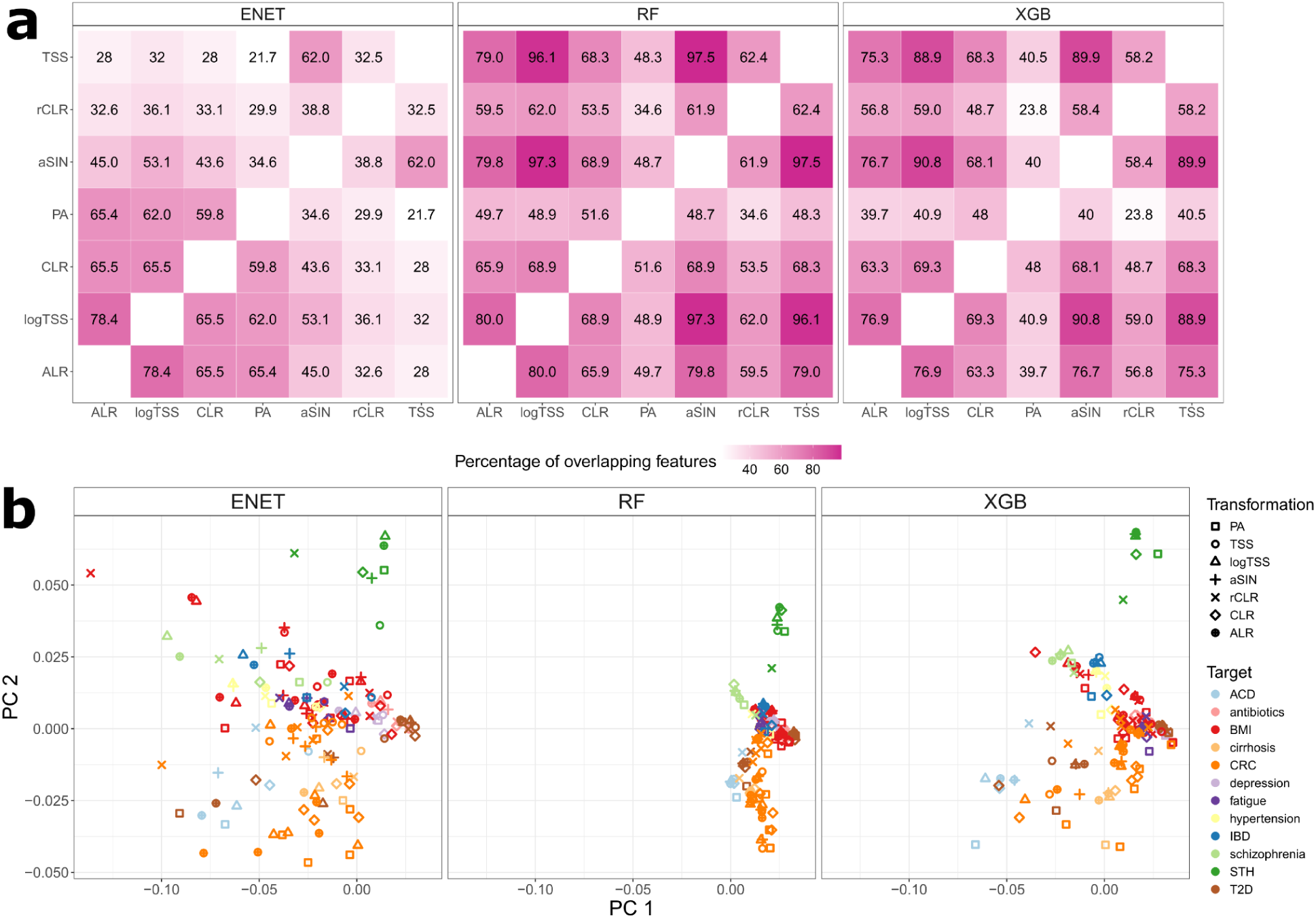
STABILITY OF FEATURE IMPORTANCE RESULTS. **a** - Overlap of top-25 features between transformations for each algorithm and dataset. **b -** Principal component analysis of the SHAP profiles visualized separately for ENET, XGB and RF. **c** - Pearson correlation between feature’s mean absolute SHAP value, prevalence, and abundance for each transformation and every algorithm. Abbreviation: BMI - body mass index; ACD - atherosclerotic cardiovascular disease; CRC - colorectal cancer; IBD - irritable bowel disease; STH - soil-transmitted helminths; T2D - type 2 diabetes.

Interestingly, although the classification performance of PA with RF was comparable or superior to CLR, rCLR, ALR and logTSS, the overlap among the most significant predictors is lowest for PA with around 50% correspondence to the predictors informed by the abundance-based data transformations. Thus, different abundance-based transformations can inform highly similar biomarker candidates, but the selection might not be optimal for classification performance indicating that PA transformation is able to indicate potentially novel biomarkers of equal predictive value. Following the observation of poor prediction performance of ENET in combination with TSS and rCLR, the informed features are also highly distinct from predictors informed by the other data transformations. Thus, our results raise caution against biomarker detection using ENET in combination with TSS. When comparing the predictors informed by RF and ENET, around 50% of the predictors overlap across the transformations. Here, PA is a notable exception with significantly higher overlap between the features informed by RF and ENET. Interestingly, RF and XGB showed a high percentage of overlapping features regardless of the data transformation used, with PA (presence-absence) having the lowest overlap. This lower overlap might stem from the fact that other transformations tend to identify a larger number of important features compared to the presence-absence data transformation (**Supplementary Figure 6)**.

Next, we assessed how similar in general are the feature importance profiles informed by different data transformations and algorithms. For that, we calculated Euclidean distances on the SHAP value profiles and carried out principal component analysis (**Figure 4b**). The results show that the target-specific signatures are clearly evident independent of the learning algorithm and data transformation. For example, predictive signatures for colorectal cancer (orange) and soil-transmitted helminths (green) clearly stand out along the PC2 axis. However, there is still a remarkable difference between the algorithms with RF displaying less variation in the feature importance profiles compared to ENET and XGB. Although feature importance profiles resulting from different data transformations are more similar within the same study than they are between studies indicating a detected target-specific signal, the effect is stronger for RF (**Supplementary Figure 7)**. Thus, RF identifies a more target-specific signal and is less affected by the choice of the data transformation. Interestingly, the feature importance profiles for ENET and XGB are more similar across studies when the same data transformation is used than they are across studies and across different data transformations (p = 1.4e-05 for ENET and p = 0.0092 for XGB). Thus, ENET and XGB can identify signatures which are not only specific to the target variable but are common across the datasets. The same is not true for RF (p = 0.17).

Given the differences in the most informative predictors, particularly between PA and abundance-based transformations, we were interested in whether the feature selection is associated with the prevalence and relative abundance of the microbial taxa. For that, we calculated Pearson correlations between SHAP values and taxa prevalence and relative abundance (**Supplementary Figure 8)**. Expectedly, we found that the SHAP values are positively correlated with the taxa prevalence. However, the correlation with prevalence was lower for the features informed by PA and CLR independent of the learning algorithm. Similarly, we observed that the SHAP values obtained by abundance-based transformations exhibit remarkably higher correlations with the relative abundance as compared to SHAP values obtained by PA. This can partially be the reason why ENET and XGB choose similar predictors across different target variables. To further understand how this correlation is reflected in the feature selection, we analyzed the microbial predictors, which had the largest difference in SHAP values between TSS and PA (**Supplementary Figures 9, 10, 11**). For all algorithms, more abundant taxa have higher significance with TSS when compared to PA. Moreover, the bacteria which were more important with the TSS include well-known and abundant gut commensals such as *Prevotella copri (Gacesa et al. 2022)*, *Bacteroides vulgatus (Lin et al. 2023), Bacteroides uniformis (Zafar and Saier 2021)*, and *Faecalibacterium prausnitzii (Lopez-Siles et al. 2017)*. Bacteria having the highest SHAP values when using any model with PA were probiotic candidates such as *Akkermansia municiphila (Cani et al. 2022)* and potantially beneficial microbes such as *Firmicutes bacterium CAG:95* and *Firmicutes bacterium CAG:110* (Zeybel et al. 2022). Models using PA as data transformation assigned also higher SHAP values to opportunistic pathobionts such as *Desulfovibrio piger (Singh, Carroll-Portillo, and Lin 2023)*, *Fusobacterium nucleatum (Brennan and Garrett 2019)* and *Erysipelatoclostridium ramosum* (previously known as *Clostridium ramosum*) (Bojović et al. 2020). This observation further highlights the difference in the biomarker profiles informed by different data transformations. With abundance-based transformations we are more likely to identify more prevalent and abundant taxa which might potentially complicate the identification of disease-specific markers. In contrast, the biomarkers informed by PA are largely independent of these properties.

In conclusion, our findings underscore that models assign high importance to a limited set of microbes, a property common for all data transformations. However, despite obtaining similar classification performance, there can be large differences in the most informative features chosen by the transformations. This variability could impact the development of gut microbiome health indices, assessment of microbiome dysbiosis, and biomarker discovery. Together with the equivalency in the resulting classification performance, this highlights the need to improve the feature identification, validation and stability.

## Discussion

Our goal was to evaluate the impact of data transformations on machine learning performance in microbiome binary classification (Marcos-Zambrano et al. 2021). We compared the classification performance of two learning algorithms in combination with seven data transformations across various analytical scenarios and analyzed the impact of using different data transformations on the feature importance. Our results showed that there was no significant improvement in classification performance when abundance-based transformations were used instead of presence-absence (PA). This result is consistent with the comparison of PA and TSS as reported by Giliberti *et al*. (Giliberti et al. 2022), but we further extend their findings to several other commonly used data transformations. Similarly to Giliberti *et al*., we observed that an elastic net algorithm (ENET) performs better with PA when compared to total-sum scaling (TSS), but there is no major difference when random forest (RF) or XGBoost (XGB) are used. Interestingly, all three models showed a decline in performance when applied with the isometric log-ratio (ILR) transformation, with the most significant drop observed in ENET. Although we saw the previously reported improved performance of ENET in combination with CLR as compared to TSS (Quinn and Erb 2020), our results didn’t confirm the benefit of using abundance-based data transformations as reported in several other studies. For example, in combination with RF, centered log-ratio (CLR) has been shown to outperform TSS and logTSS (Kubinski et al. 2022). Moreover, based on our results there was no significant effect of the transformation selection on the model generalizability nor interaction with the model performance and data dimensionality. The performance of machine learning models seems to be more reliant on the characteristics of the dataset than on the specific algorithm or transformation employed. This principle echoes the ‘no free lunch theorem’ in machine learning, emphasizing that no single algorithm universally outperforms others across diverse datasets (Wolpert and Macready 1997).

Our feature analysis showed that the number of significant predictors informed by the model was more tied to the dataset than the transformations used. Importantly, there was only partial overlap among the top features informed by different data transformation, indicating that different sets of microbial features can yield similar classification performance. This observation was most pronounced for PA in combination with RF which had the lowest overlap with the other data transformations, although PA led to better predictive performance. Thus, microbiome studies could take advantage of the methodologies such as statistically equivalent signatures, which aim to identify variable sets with equal predictive power (Lagani et al. 2017). Analyzing the properties of the selected features shows that abundance-based transformations may focus on the most abundant gut microbes such as *P. copri*, *F. prausnitzii, B. vulgatus* and *B. uniformis*. As using relative abundances as a data representation is one of the most popular choices for applying ML on the microbiome data (Marcos-Zambrano et al. 2021), we suggest taking caution when focusing on the feature importance as the selection might not yield a signal specific to the condition of interest and it could potentially obscure the interpretation of classification outcomes. These findings underscore the need for further research.

Based on our findings, we recommend adopting the presence-absence (PA) transformation for microbiome data classification tasks as a supportive alternative to the abundance-based transformations. PA-based classifiers demonstrate strong performance, offer a simpler interpretation, avoiding the need for pseudocount imputation or data scale transformation. However, selecting a threshold for microbe absence (e.g., setting it at zero) introduces challenges related to structural zeros and sequencing depth, which warrants further investigation (Silverman et al. 2020).

Our study’s strengths lie in its systematic approach, employing two learning algorithms on diverse datasets, which enhances generalizability. However, we acknowledge limitations. We recognize that other data transformations, such as PhyILR (Silverman et al. 2017), may outperform the ones we used. However, our focus was on the most commonly used transformations in microbiome studies to provide a solid starting point for researchers interested in applying machine learning to binary classification tasks. Our focus is on classification tasks, potentially limiting relevance to other analyses. Dataset constraints and unexplored confounding variables are also noted. To further understand the effect of data transformation on classification accuracy and feature selection, (Gaulke and Sharpton 2018)taking advantage of synthetic microbial communities where alterations in bacteria are documented and ground truth is available would be beneficial. Such an approach is being used for example *in silico* gut microbiome community design (Baranwal et al. 2022; Venturelli et al. 2018; Clark et al. 2021).

In the future, we plan to explore additional questions. Firstly, we would like to assess whether using only presence-absence (PA) data is sufficient for accurate classification or if combining PA with variations in the abundance of key bacteria enhances performance. We would also investigate if the observed similarity between PA-based classifiers and abundance-based transformations is influenced by technical factors, such as shallow sequencing, which might mask certain features. Additionally, we aim to gather more datasets to assess whether SHAP values for bacterial features in each disease remain consistent across geographic locations. Geographic location is a known confounder in microbiome studies due to the significant influence of regional diets on gut microbiome composition. Moreover we would like to extend our study and evaluate how data transformations impact various types of data modeling tasks, including regression models and unsupervised techniques like clustering. These inquiries highlight the complexity of our findings and provide directions for further research.

## Methods

### Data acquisition

#### Open data

Datasets available in the *curatedMetagenomicData* R package were used for the analysis (version 3.6.2) (Pasolli et al. 2017). The *curatedMetagenomicData* package contains uniformly processed shotgun metagenomic sequencing human microbiome data of healthy and diseased subjects. The microbiome data preprocessing, including taxonomic profiling, is carried out using the bioBakery 3 toolkit *(Beghini et al. 2021)*. For the within-study (WS) evaluation setting, we focused on 17 distinct human stool metagenomics datasets, which had a primary endpoint available and included at least 50 cases and 50 controls to allow proper model evaluation. Also, we included four datasets where we defined obesity as a binary outcome defined as BMI > 30. In some instances, filtering was applied to the original dataset to achieve a binary classification task. For example, we excluded adenomas from the Zeller *et al*. study and focused only on colorectal cancer cases and controls (Zeller et al. 2014). The selected datasets, sample size, defined binary classification task and the filtering procedures are shown in **Supplementary Table 1**.

For the leave-one-study-out (LOSO) evaluation setting, we included additional 6 colorectal cancer datasets with less than 50 cases or controls and 4 datasets, where primary endpoint was not available.

#### Estonian microbiome cohort

The Estonian Microbiome cohort (EstMB) is a volunteer based cohort currently including 2509 subjects, who have provided blood, oral and stool samples. Being part of a larger nation-wide Estonian Biobank (EstBB), linkings to various electronic health records (EHR) and questionnaires covering the lifestyle and dietary preferences are available for all of the subjects. The cohort is described in detail in (Aasmets et al. 2022). For the binary classification, four target variables were considered: antibiotics usage within the previous 90 days before the microbiome sample collection (AB90), obesity defined as BMI > 30 (BMI30), type 2 diabetes (T2D) and depression (**Supplementary Table 1)**.

Taxonomic profiling on EstMB was carried out using *Metaphlan3 (Beghini et al. 2021)* to comply with the profiling done for the *curatedMetagenomicData* R package datasets.

### Data transformations

Numerous data transformations have been proposed to be used in the analysis of microbiome data. In the current manuscript, the following data transformations were considered:

1. Relative abundance/total-sum-scaling (TSS): The standard and most widely used technique, which scales data to relative abundances.
2. Log(TSS): A logarithmic transformation applied to TSS-normalized data.
3. Presence-absence (PA): changes abundance data into binary data. We used zero as the threshold for presence-absence.
4. Arcsin square root (aSIN (Delgado et al. 2019)): Which involves applying the arcsin (inverse sine) function to each value, which can be useful for normalizing and stabilizing data that represents proportions or percentages, particularly in statistical analyses of compositional data.
5. Centered log-ratio transformation (CLR (Aitchison 1982)): procedure that enhances compositional data for standard statistical analysis by dividing each value by the geometric mean of all features and applying a logarithmic transformation.
6. Robust CLR transformation (rCLR (Martino et al. 2019)): a robust version of CLR that handles zeros by using only observed taxa for the geometric mean calculations.
7. Additive log-ratio transformation (ALR (Aitchison 1982)): in which each feature in a dataset is divided by a selected reference feature and then logarithmically transformed. We randomly selected 5 features as reference elements and averaged all the results over the different ALR transformations to account for the variability arising from the reference element selection. We observed that averaging over 5 different reference elements resulted in a reasonable trade-off between computational burden and variability of the performance estimates **(Supplementary Figure 12).**
8. Isometric log-ratio transformation (ILR <u>(Egozcue et al. 2003)</u>): in which the compositional dataset is transformed by representing each feature as a set of orthogonal log-ratios using a basis that maintains the geometric structure of the data. The ILR transformation was applied using the implementation in the R package *compositions*.

### Machine learning pipeline

Each of the considered transformations was applied to a dataset and a binary classification task was carried out. Random forest (RF), XGBoost (XGB)and elastic net (ENET) penalized logistic regression were used as the learning algorithms. ENET logistic regression is a machine learning algorithm that combines L1 (Lasso) and L2 (Ridge) regularization techniques to perform logistic regression with variable selection, making it suitable for high-dimensional data by minimizing overfitting and selecting the most relevant features. Regularization helps prevent overfitting by adding a penalty to the model’s complexity. L1 regularization (Lasso) encourages sparsity by shrinking some weights to zero, which helps feature selection. L2 regularization (Ridge) distributes penalties evenly across all weights, reducing their magnitude but keeping all features. A balance between L1 and L2 (Elastic Net) combines these benefits, offering both feature selection and weight regularization, helping the model generalize better. (Zou and Hastie 2005). Random Forest, on the other hand, is an ensemble learning method that builds multiple decision trees and combines their predictions to improve classification accuracy and handle complex relationships in data while reducing the risk of overfitting (Breiman 2001). XGBoost, is an advanced ensemble learning algorithm that builds multiple decision trees sequentially, where each tree corrects the errors of the previous ones, thereby improving the overall prediction accuracy. It is designed to be highly efficient and scalable, handling large datasets with complex relationships while preventing overfitting through regularization (Chen and Guestrin 2016). For every algorithm we performed hyperparameter tuning using cross validation in combination with grid search and the model with optimal hyperparameters was then used for classification. For ENET, both penalty and mixture parameters were tune; for RF the number of predictors to sample at each split (*mtry*) and the number of observations needed to keep splitting nodes (*min_n*) were tuned, the number of trees was 500; for XGB, for RF the number of predictors to sample at each split (*mtry*) and the number of observations needed to keep splitting nodes (*min_n*), the maximum depth of each tree (*tree_depth*), learning rate, the fraction of training data used for growing each tree (*sample_size*) were tuned, the number of trees was 500.

These algorithms were chosen due to their popularity and competitive performance in the microbiome field (Zhou and Gallins 2019) and as RF being a nonlinear, XGB being non-linear and using boosting, where trees are built sequentially, with each tree focusing on correcting the mistakes of the previous ones and ENET a linear method, they can provide insights on the impact of algorithm selection in microbiome studies. Followingly, the model fitting and evaluation is described.

#### Within the study (WS) setting

For each 21 datasets used for the within-study evaluation, the following repeated holdout validation procedure for parameter tuning and model evaluation was carried out:

1. *Data is split to training/test set (75%-25%) stratified by the target variable*
2. *Hyperparameter tuning on the training set (75%) using 5-fold cross-validation with grid search (10 parameter combinations)*
3. *Model with optimal hyperparameters is fit on the whole training data (75%)*
4. *Model is evaluated on the test set (25%)*

The initial data test/train split and model evaluation were carried out on 10 random data splits to assess the variation arising from sampling resulting in 10 performance estimates per evaluation.

#### Leave-One-Study-Out (LOSO) setting

The LOSO setting was carried out for the 11 available colorectal cancer and 5 obesity (BMI >= 30) datasets. The aim was to understand whether the dataset-to-dataset generalization performance might be dependent on the chosen data transformation. For the model fitting and evaluation, the following procedure was carried out:

1. *Data is split to training/test set so that one dataset works as the test set and other datasets as a combined training set*
2. *Hyperparameter tuning on the training set using 5-fold cross-validation with grid search (10 parameter combinations)*
3. *Model with optimal hyperparameters is fit on the whole training data (75%)*
4. *Model is evaluated on the test set - left out dataset*

The model evaluation was carried out using each dataset per target variable as a test set. This resulted in 11 performance estimates for colorectal cancer and 5 for obesity.

#### Cumulative classifier

This analysis aimed to evaluate whether a subset of the most significant predictors can be used to build a model that has optimal prediction performance. This experiment was based on the PA-transformed EstMB datasets and carried out for antibiotics usage, obesity (BMI30) and depression. For the model fitting, feature selection and evaluation, the following procedure was carried out:

*Stage 1*

1 *Data is split to training/test set (50%-50%) stratified by the target variable*
2 *Hyperparameter are tuned (5-fold cross-validation with grid search) and feature importance calculated on the training set (50%)*
3 *Subsets of most important features are created (set sizes 5, 10, 25, 50, 75, 100, 200)*

*Stage 2*

4 *Test data from stage 1 (50%) is used for model evaluation*
5 *For each subset of features:*

a. *Stage 1 test data is split to stage 2 training/test set (75%-25%) stratified by the target variable*
b. *Hyperparameter tuning on the stage 2 training set (75%) using 5-fold cross-validation with grid search (10 parameter combinations)*
c. *Model with optimal hyperparameters is fit on the whole stage 2 training data (75%)*
d. *Model is evaluated on the stage 2 test set (25%)*

The stage 1 and stage 2 data test/train splits and model evaluation were carried out on 10 random data splits to assess the variation arising from sampling.

### Model comparison

To test the differences in the binary classification results, we used the Wilcoxon signed-rank test for each pair of data transformations. For that, we first averaged the results over different folds and target variables to account for the overrepresentation of certain phenotypes like colorectal cancer and obesity. After that, a paired test within each target variable was carried out. Thus, we tested the hypothesis that the difference in the performance of *transformation1* and *transformation2* is not equal to 0 (**Figure 1c**). To account for the multiple testing, Benjamini-Hochbergi procedure was applied to the nominal p-values.

### Filtering subjects and features

Due to the large sample size, EstMB dataset was further used to study the effects of sample size and number of bacteria used by the model on the performance of the classifiers, focusing on differences between the data transformations. For that reason, the cases and controls of antibiotics usage (AB90) and obesity (BMI30) were subsampled to 20/40/60/80% of the initial number of cases and controls. Additionally, in combination with the sample subsetting, prevalence filtering (1/10/50%) for the microbial taxa was applied to study the impact of the number of predictors. Thus, in total of 5*3 = 15scenarios per algorithm and data transformation were analyzed and the same machine learning procedure as described in the within-sample setting evaluation was carried out.

Prevalence filtering (10/25/50/75/90%) was also carried out on the *curatedMetagenomicData* package datasets before applying the machine learning procedure as described in the within-sample setting evaluation was carried out.

### Feature importance analysis

Feature importance evaluation was based on the SHapley Additive exPlanations (SHAP) values. SHAP values are a method used in machine learning to explain the contribution of each feature to the prediction of a model (Lundberg and Lee 2017). We first determined the percentage of features within each dataset that exhibited mean absolute SHAP values exceeding zero. For feature overlap assessment, we quantified the average percentage of overlapping features among the top 25 features between data transformations and algorithms. The average feature overlap was calculated by taking the average across different folds and datasets. Pearson correlation was used to study the associations between SHAP values, bacterial relative abundances and prevalence. Overall similarity of the feature importance profiles was evaluated by comparing the Euclidean distances between the feature importance profiles across the algorithm-transformation pairs. Standard t-test was used to formally test the differences between groups of interest (**Supplementary Figure 7)**. Principal component analysis on the feature importance profiles was carried out to visualize the differences between different data transformations and datasets in a two-dimensional space.

To identify microbial predictors with the largest difference in SHAP values between TSS and PA we first calculated the mean absolute SHAP value for each feature, study, data transformation and algorithm. Then, for each feature, study and algorithm we calculated the delta between the mean SHAP values for TSS and PA and averaged the deltas over all studies. Top 100 features according to the absolute delta for algorithms separately were then visualized (**Supplementary Figures 9, 10, 11**).

## Code availability

The source code for the analyses is available at https://github.com/bioinf-mcb/transformation_benchmark.

## Acknowledgments

The authors would like to thank Triinu Temberg, Mari-Liis Tammeorg, Marili Palover, Anu Reigo, Neeme Tõnisson, Liis Leitsalu, and Esta Pintsaar for participating in the sample collection process of the Estonian Microbiome cohort. We thank Steven Smit, Rita Kreevan and Martin Tootsi, for the DNA extraction process. We thank Reidar Andreson for bioinformatic support and Witold Wydmański for fruitful discussions during project planning. We also thank all the study participants. This study was funded by the European Union through the European Regional Development Fund Project No. 2014-2020.4.01.15-0012 GENTRANSMED. Data analysis was carried out in part in the High-Performance Computing Center of University of Tartu.

## Conflict of Interest

The authors declare that the research was conducted in the absence of any commercial or financial relationships that could be construed as a potential conflict of interest.

## Author Contributions

E.O. and T.K., designed and supervised the study. Z.K. and O.A. performed the data analysis, interpreted the data and prepared the figures. Z.K. and O.A. wrote the manuscript. All authors read and approved the final paper.

## Funding

This work was funded by an Estonian Research Council grant (PRG1414 to EO) and an EMBO Installation grant (No. 3573 to EO). TK and ZK are funded by the National Science Centre, Poland grant 2019/35/D/NZ2/04353.

## Supplementary Figures

**Supplementary Figure 1.**
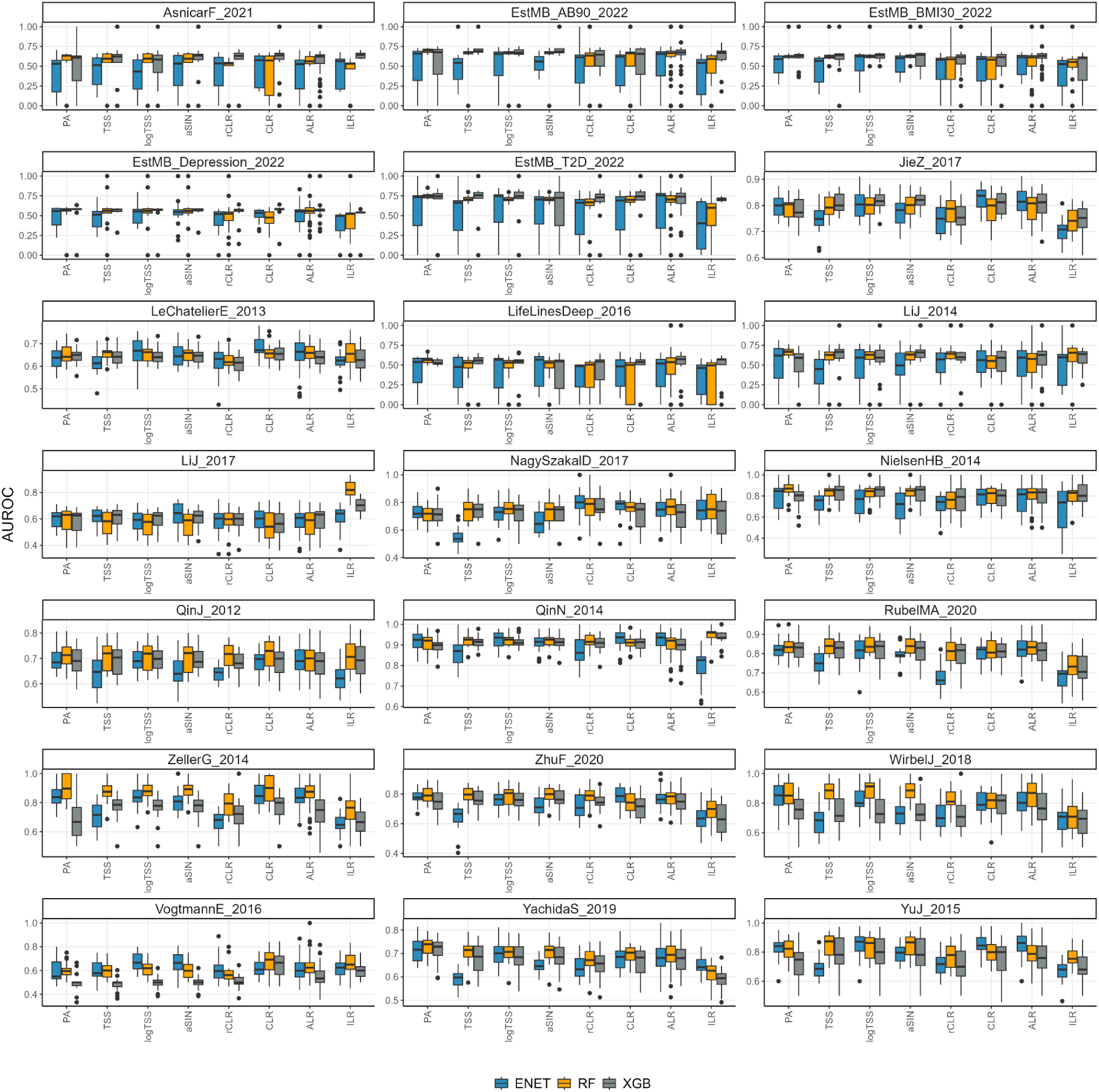
Classification performance (AUROC) by studies. Abbreviations: *ACD - atherosclerotic cardiovascular disease;* BMI30 *- body mass index > 30; CRC - colorectal cancer; IBD - inflammatory bowel disease; STH - soil-transmitted helminths*; *T2D - type 2 diabetes*

**Supplementary Figure 2.**
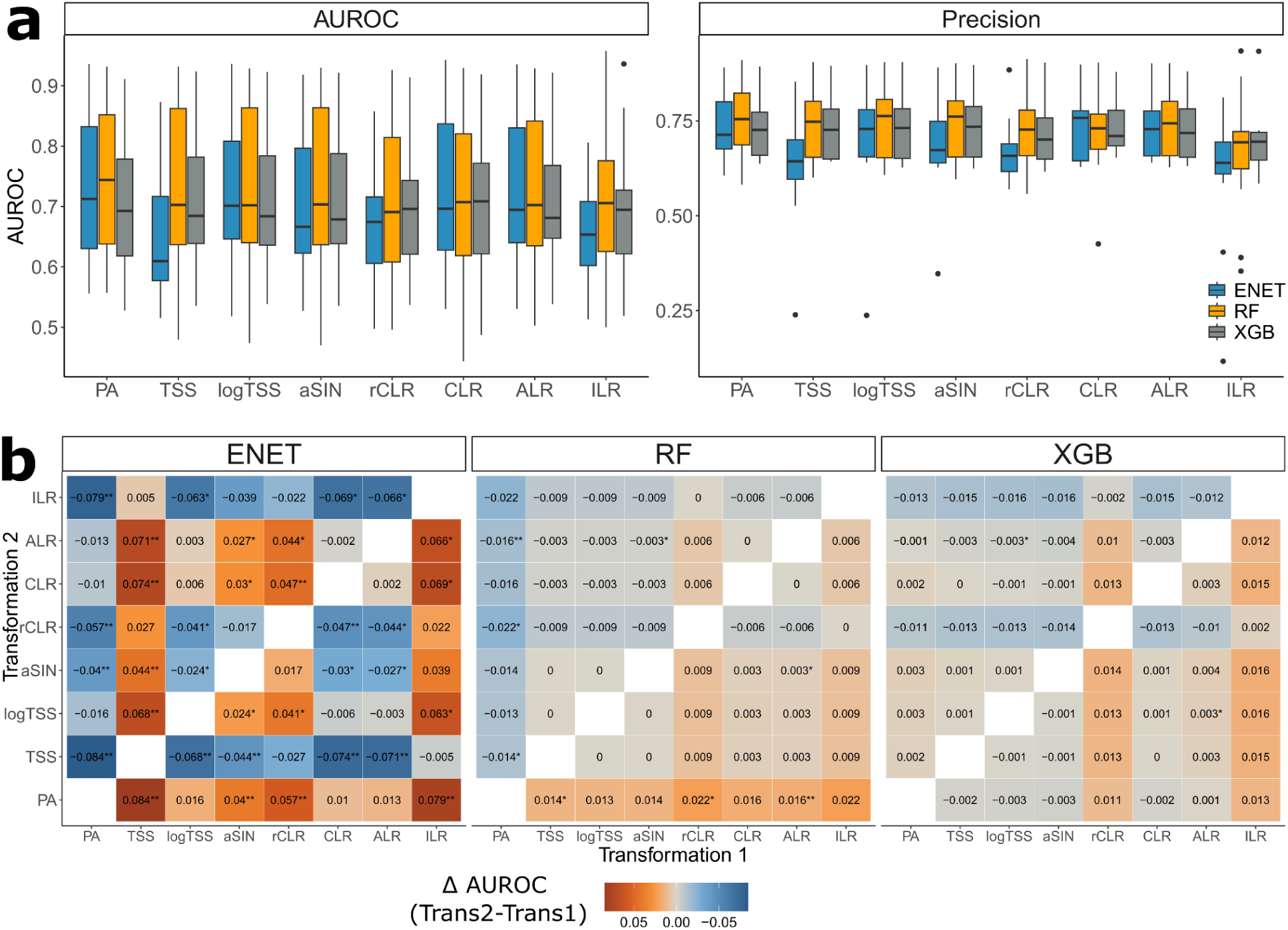
**a** - Classification performance (AUROC and precision) in the within-study setting for every data transformation and algorithm for the rarefied data. **b -** Statistical analysis results between the used data transformations for rarefied data (Wilcoxon signed-rank testpaired t-test) for elastic net (ENET), and random forest (RF) and extreme gradient boosting (XGB); Values and colors correspond to the differences in AUROC between Transformation 2 and Transformation 1; * indicates a nominally statistically significant difference in AUROC (Wilcoxon signed-rank test, p-value ≤ 0.05), ** indicates a statistically significant difference in AUROC after correction (Wilcoxon signed-rank test, FDR ≤ 0.05). Abbreviations: ENET - elastic net logistic regression; RF - random forest; XGB - extreme gradient boosting, XGBoost; PA - presence-absence; TSS - total-sum scaling; logTSS - logarithm of TSS; aSIN - arcsine square root; CLR - centered log-ratio; rCLR - robust CLR; ALR - additive log-ratio; ILR - isometric log-ratio.

**Supplementary Figure 3.**
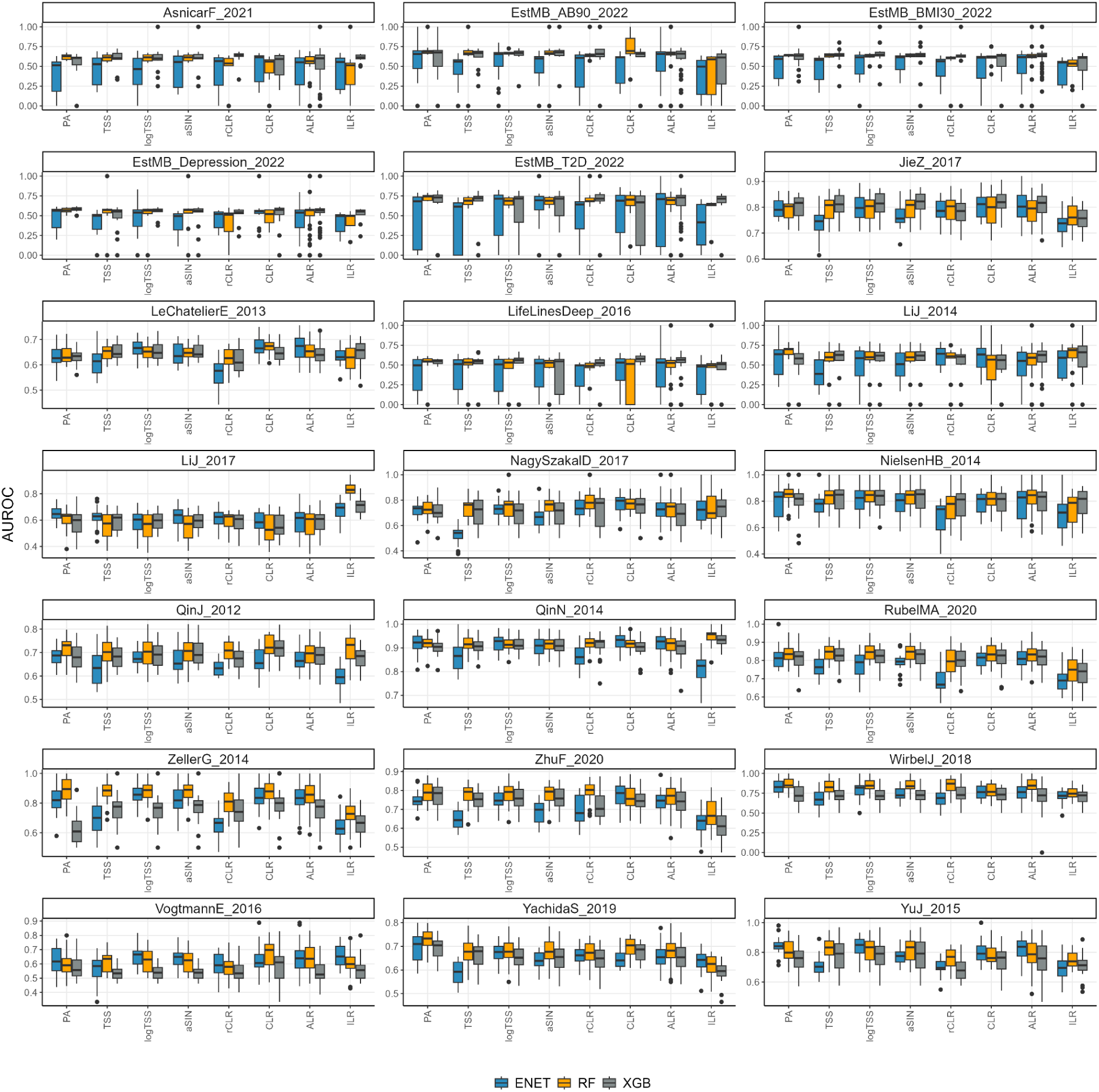
Classification performance (AUROC) by studies on rarefied data. Abbreviations: *ACD - atherosclerotic cardiovascular disease;* BMI30 *- body mass index > 30; CRC - colorectal cancer; IBD - inflammatory bowel disease; STH - soil-transmitted helminths*; *T2D - type 2 diabetes*

**Supplementary Figure 4.**
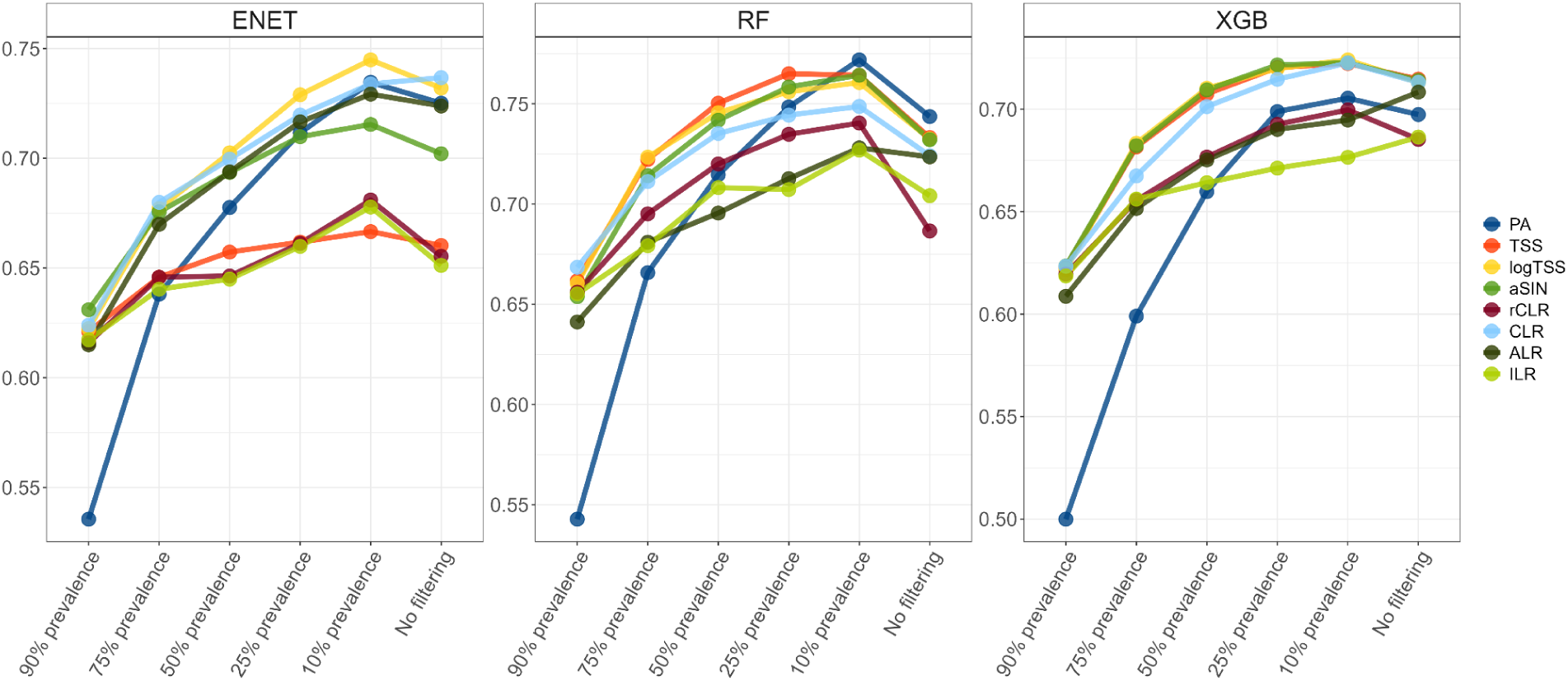
Classifier performance dependence on the data transformation and data dimensionality (feature filtering based on the prevalence of the taxa). Colors represent data transformation.

**Supplementary Figure 5.**
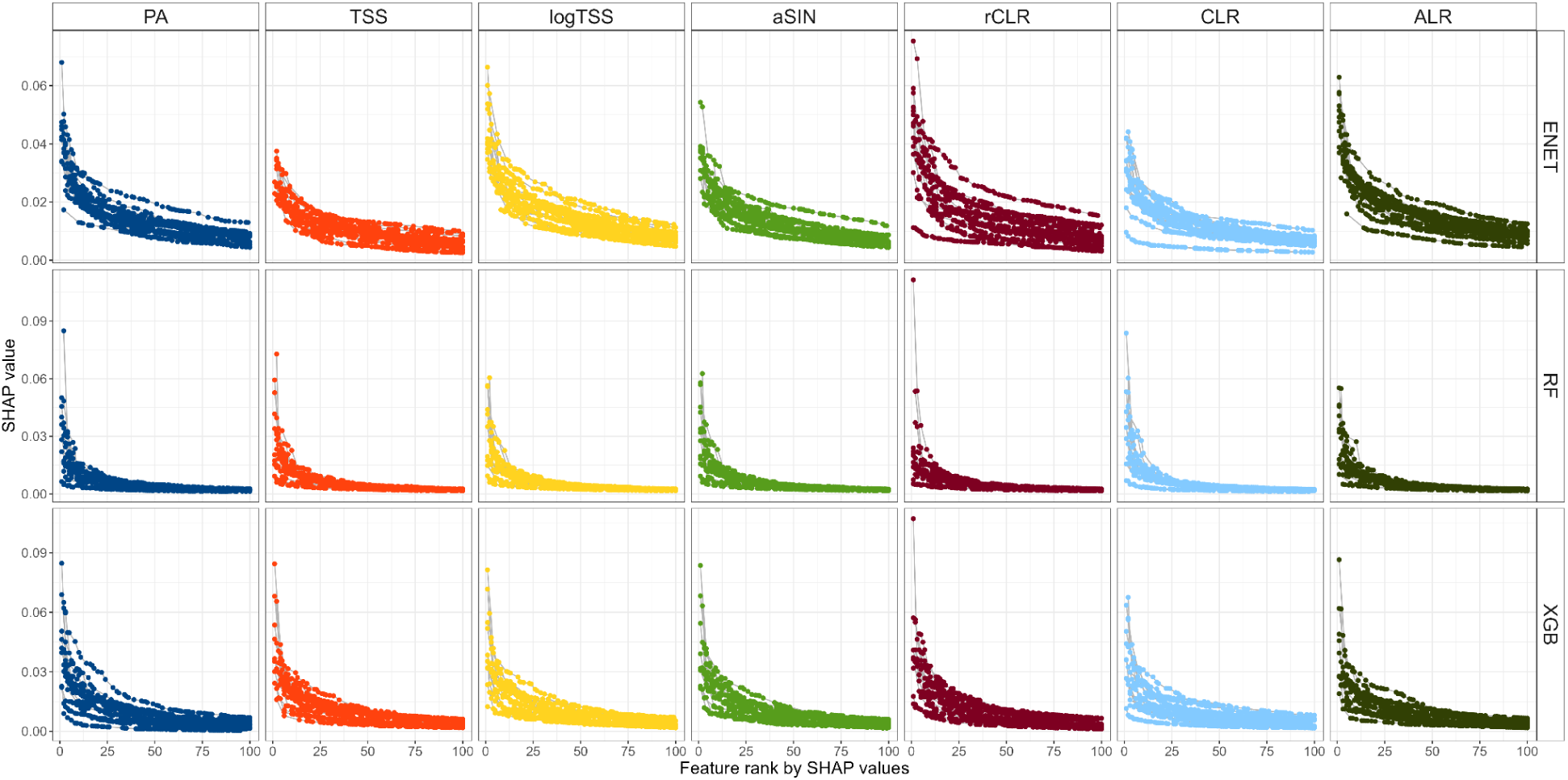
Mean absolute SHAP values distribution for top 50 features across all datasets plotted separately for each transformation.

**Supplementary Figure 6.**
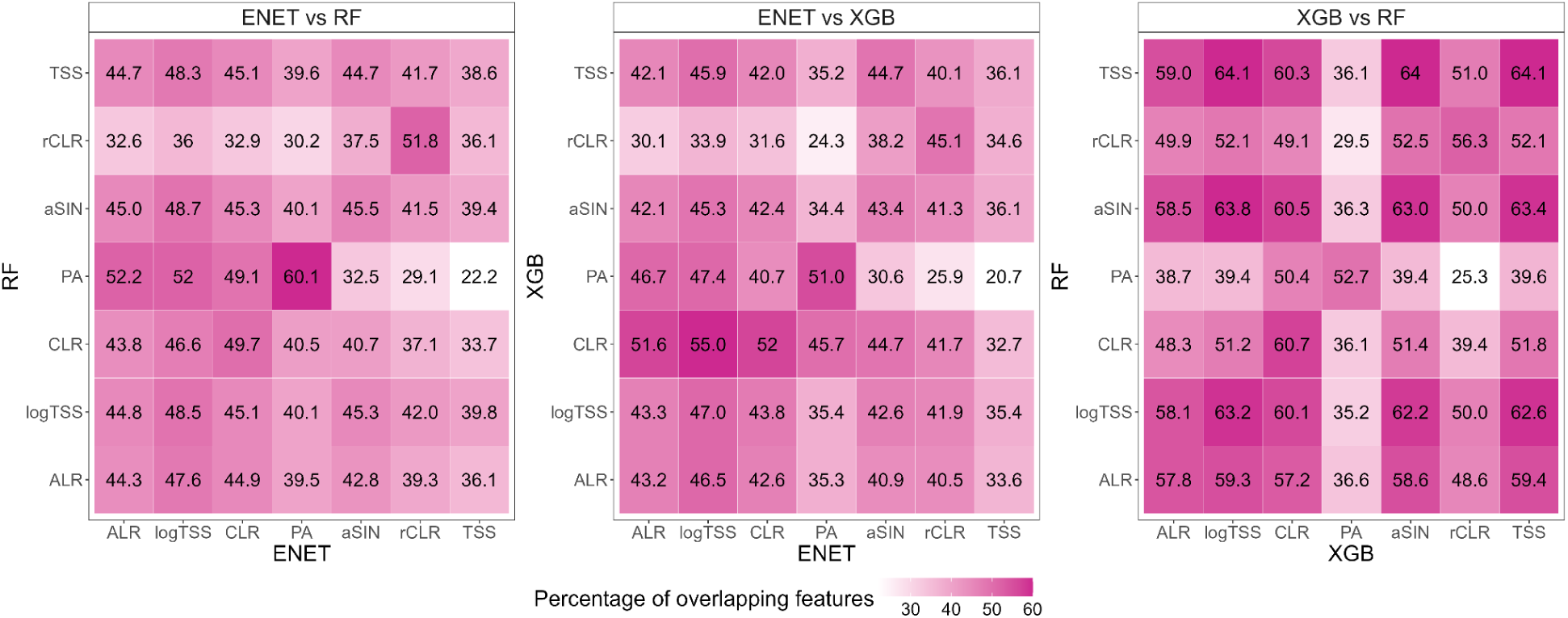
Overlap in the most significant predictors (top 25 predictors by SHAP values) between the algorithm-data transformation combinations.

**Supplementary Figure 7.**
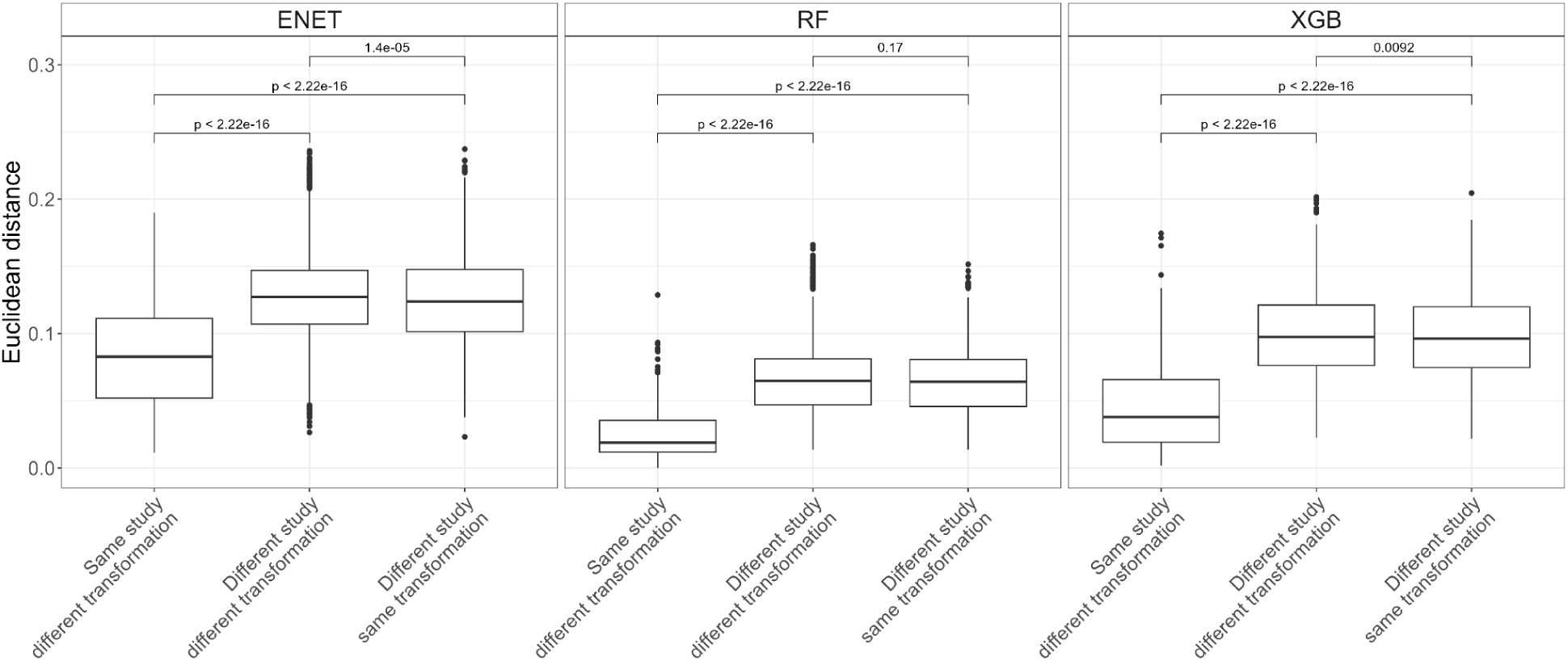
Differences in the feature importance profiles between and within different studies and data transformations measured by Euclidean distance. P-values correspond to the unpaired t-test.

**Supplementary Figure 8.**
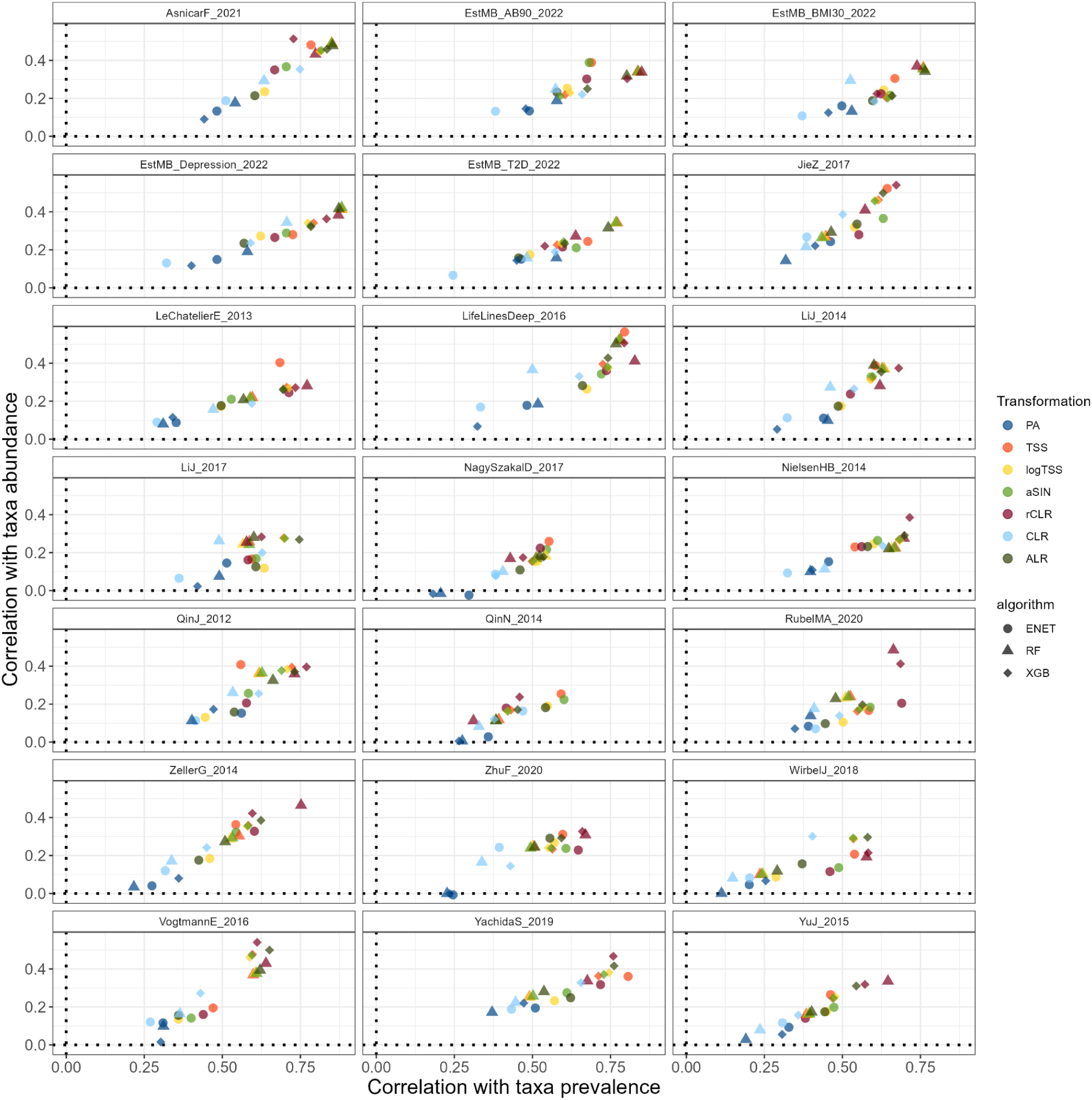
Correlation between the mean absolute SHAP values and taxa prevalence/mean relative abundance. Abbreviations: *ACD - atherosclerotic cardiovascular disease; BMI - body mass index; CRC - colorectal cancer; IBD - inflammatory bowel disease; STH - soil-transmitted helminths*; *T2D - type 2 diabetes*

**Supplementary Figure 9.**
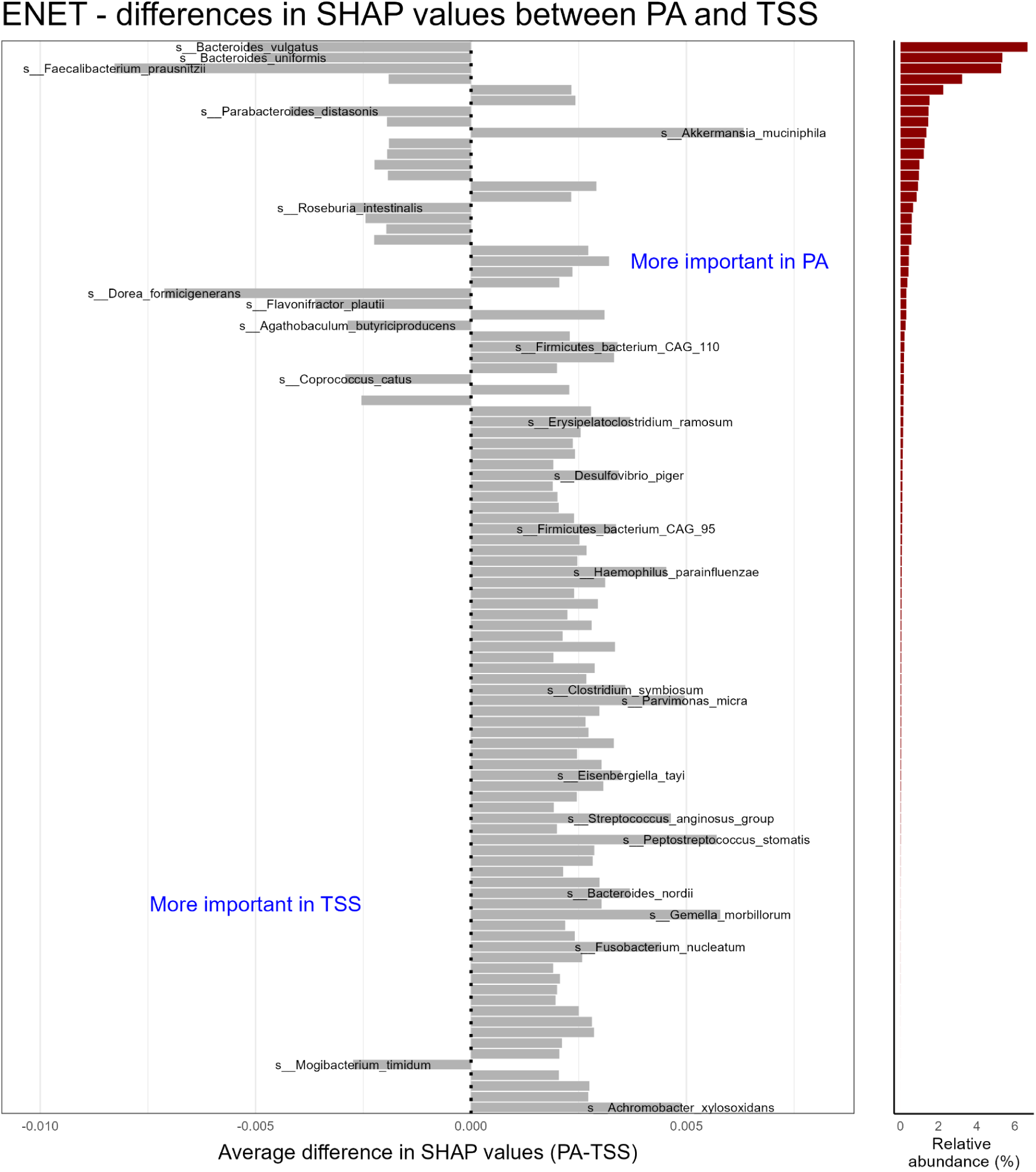
Mean differences in SHAP values across all datasets between PA and TSS for elastic net logistic regression (ENET). Top 100 taxa according to the absolute mean difference in SHAP values are shown. Taxa with the largest absolute mean difference in SHAP values between PA and TSS are further highlighted (z-score >= 3). Red barplots indicate the relative abundance of these taxa in percentages.

**Supplementary Figure 10.**
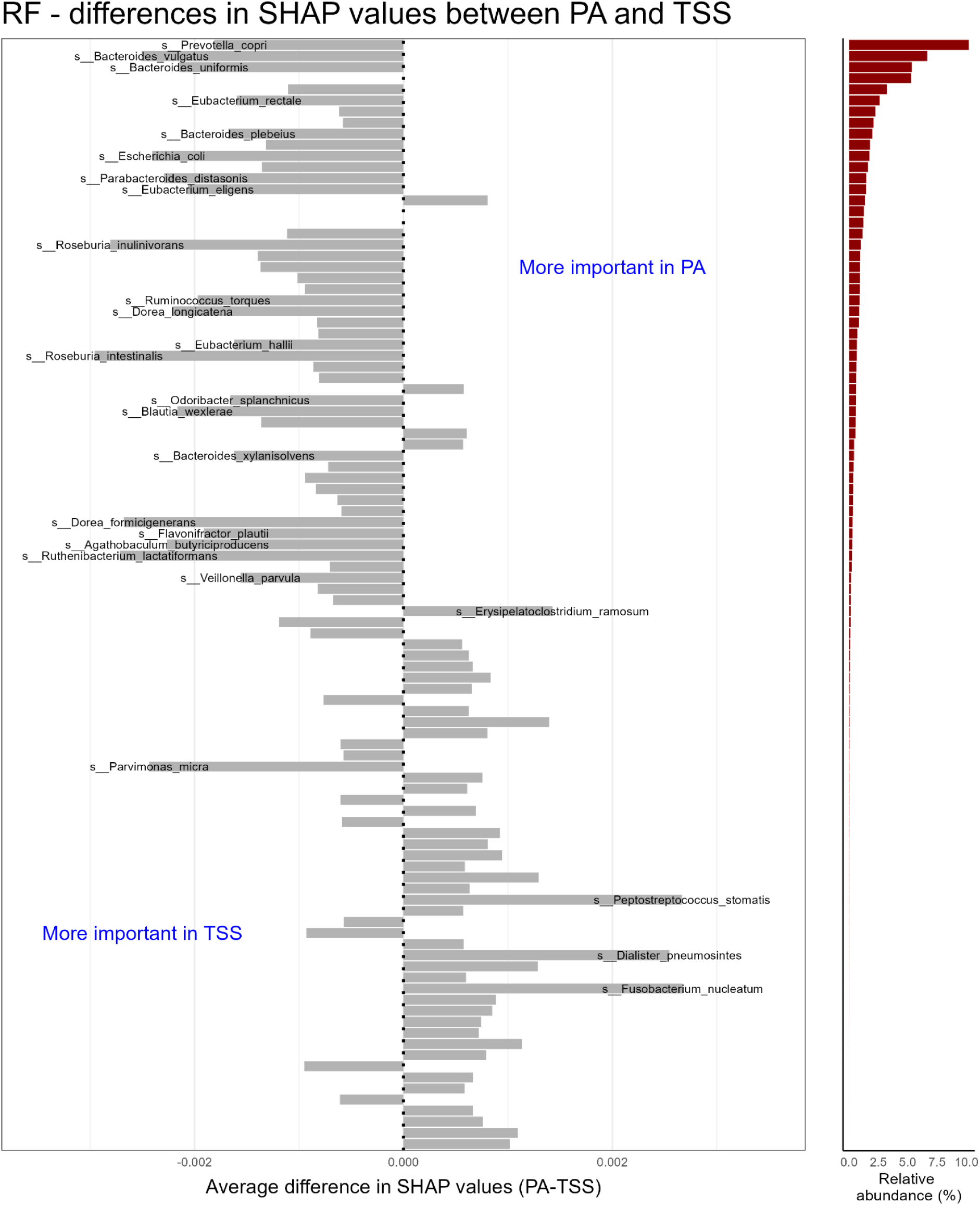
Mean differences in SHAP values across all datasets between PA and TSS for random forest (RF). Top 100 taxa according to the absolute mean difference in SHAP values are shown. Taxa with the largest absolute mean difference in SHAP values between PA and TSS are further highlighted (z-score >= 3). Red barplots indicate the relative abundance of these taxa in percentages.

**Supplementary Figure 11.**
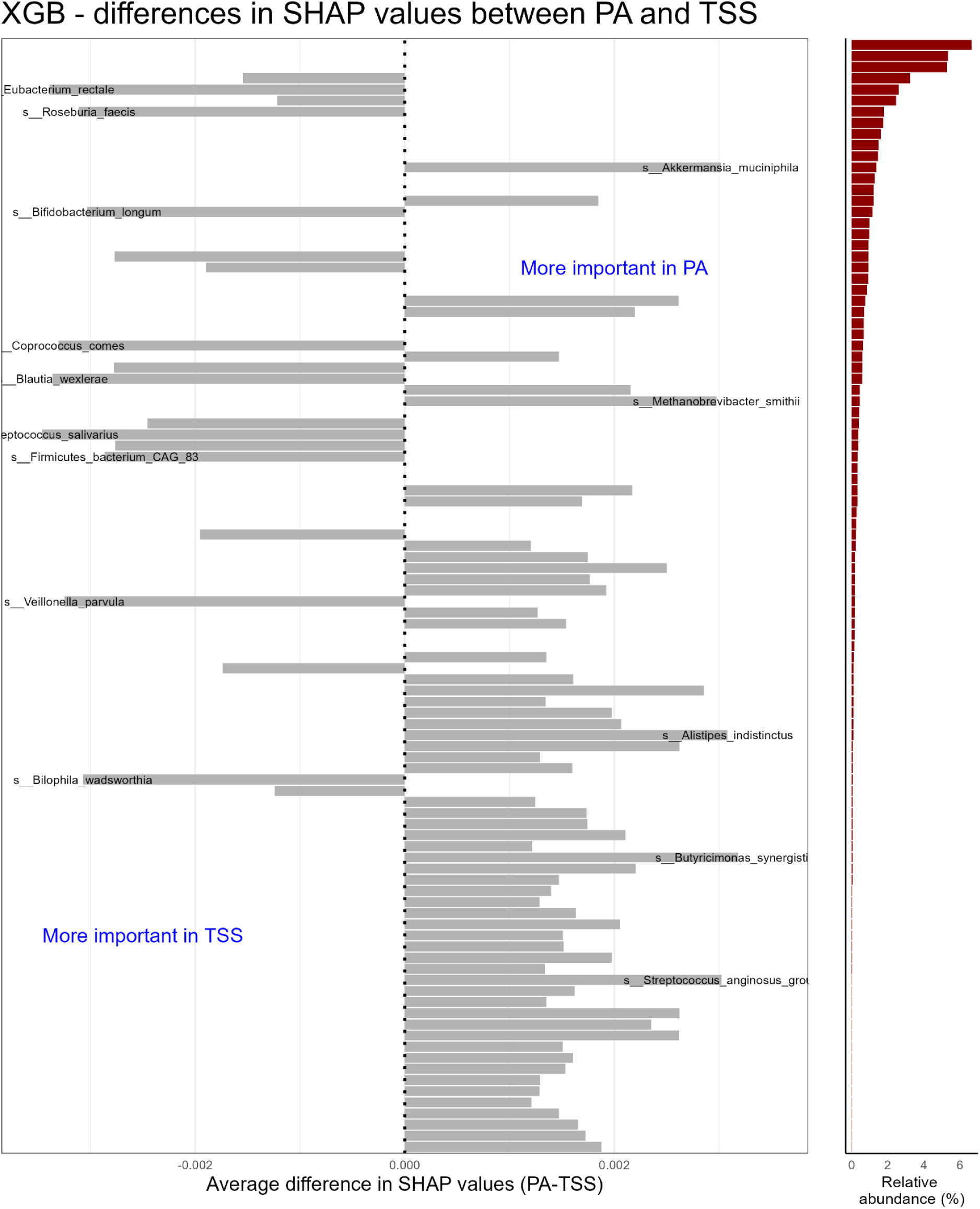
Mean differences in SHAP values across all datasets between PA and TSS for XGBoost. Top 100 taxa according to the absolute mean difference in SHAP values are shown. Taxa with the largest absolute mean difference in SHAP values between PA and TSS are further highlighted (z-score >= 3). Red barplots indicate the relative abundance of these taxa in percentages.

**Supplementary Figure 12.**
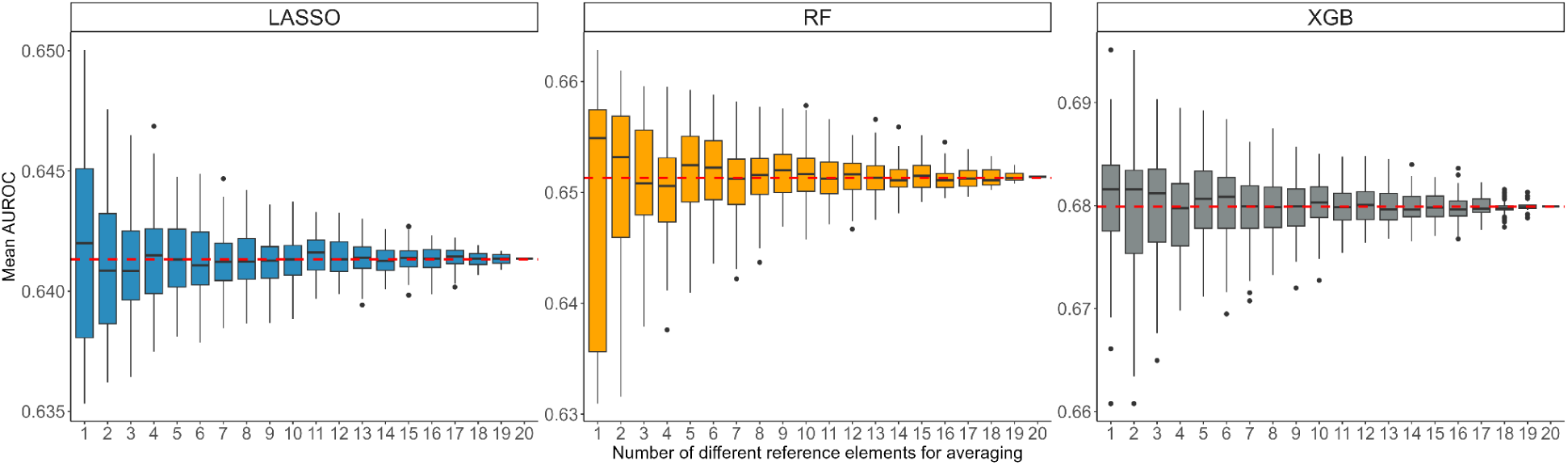
Distribution of classifier performance for antibiotics consumption in the Estonian Microbiome cohort in respect to the number of different reference elements used for averaging for the additive log-ratio (ALR) transformation.

## Notes

### Competing Interest Statement

The authors have declared no competing interest.

### Summary of Updates

In this version we updated the paper based on reviewer comments - we added XGBoost algorithm and ILR transformation.

## References

Aasmets, Oliver, Kertu Liis Krigul, Kreete Lüll, Andres Metspalu, and Elin Org. 2022. “Gut Metagenome Associations with Extensive Digital Health Data in a Volunteer-Based Estonian Microbiome Cohort.” Nature Communications 13 (1): 869.

Aitchison, J. 1982. “The Statistical Analysis of Compositional Data.” Journal of the Royal Statistical Society. Series B, Statistical Methodology 44 (2): 139–77.

Armstrong, George, Gibraan Rahman, Cameron Martino, Daniel McDonald, Antonio Gonzalez, Gal Mishne, and Rob Knight. 2022. “Applications and Comparison of Dimensionality Reduction Methods for Microbiome Data.” Frontiers in Bioinformatics 2 (February):821861.

Baranwal, Mayank, Ryan L. Clark, Jaron Thompson, Zeyu Sun, Alfred O. Hero, and Ophelia S. Venturelli. 2022. “Recurrent Neural Networks Enable Design of Multifunctional Synthetic Human Gut Microbiome Dynamics.” eLife 11 (June). 10.7554/eLife.73870.

Beghini, Francesco, Lauren J. McIver, Aitor Blanco-Míguez, Leonard Dubois, Francesco Asnicar, Sagun Maharjan, Ana Mailyan, et al. 2021. “Integrating Taxonomic, Functional, and Strain-Level Profiling of Diverse Microbial Communities with bioBakery 3.” eLife 10 (May). 10.7554/eLife.65088.

Bojović, Katarina, Ður-D Ica Ignjatović, Svetlana Soković Bajić, Danijela Vojnović Milutinović, Mirko Tomić, Nataša Golić, and Maja Tolinački. 2020. “Gut Microbiota Dysbiosis Associated With Altered Production of Short Chain Fatty Acids in Children With Neurodevelopmental Disorders.” Frontiers in Cellular and Infection Microbiology 10 (May):223.

Breiman, Leo. 2001. “Random Forests.” Machine Learning 45 (1): 5–32.

Brennan, Caitlin A., and Wendy S. Garrett. 2019. “Fusobacterium Nucleatum - Symbiont, Opportunist and Oncobacterium.” Nature Reviews. Microbiology 17 (3): 156–66.

Cani, Patrice D., Clara Depommier, Muriel Derrien, Amandine Everard, and Willem M. de Vos. 2022. “Akkermansia Muciniphila: Paradigm for next-Generation Beneficial Microorganisms.” Nature Reviews. Gastroenterology & Hepatology 19 (10): 625–37.

Chen, Tianqi, and Carlos Guestrin. 2016. “XGBoost: A Scalable Tree Boosting System,” March. 10.1145/2939672.2939785.

Clark, Ryan L., Bryce M. Connors, David M. Stevenson, Susan E. Hromada, Joshua J. Hamilton, Daniel Amador-Noguez, and Ophelia S. Venturelli. 2021. “Design of Synthetic Human Gut Microbiome Assembly and Butyrate Production.” Nature Communications 12 (1): 3254.

Delgado, Ramón Tolosana, Hassan Talebi, Mahdi Khodadadzadeh, and K. G. van den Boogaart. 2019. “On Machine Learning Algorithms and Compositional Data.” In Proceedings of the 8th International Workshop on Compositional Data Analysis (CoDaWork2019): Terrassa, 3-8 June, 2019, 172–75. Universidad Politécnica de Cataluña / Universitat Politècnica de Catalunya.

Gacesa, R., A. Kurilshikov, A. Vich Vila, T. Sinha, M. A. Y. Klaassen, L. A. Bolte, S. Andreu-Sánchez, et al. 2022. “Environmental Factors Shaping the Gut Microbiome in a Dutch Population.” Nature 604 (7907): 732–39.

Gaulke, Christopher A., and Thomas J. Sharpton. 2018. “The Influence of Ethnicity and Geography on Human Gut Microbiome Composition.” Nature Medicine 24 (10): 1495–96.

Giliberti, Renato, Sara Cavaliere, Italia Elisa Mauriello, Danilo Ercolini, and Edoardo Pasolli. 2022. “Host Phenotype Classification from Human Microbiome Data Is Mainly Driven by the Presence of Microbial Taxa.” PLoS Computational Biology 18 (4): e1010066.

Gloor, Gregory B., Jean M. Macklaim, Vera Pawlowsky-Glahn, and Juan J. Egozcue. 2017. “Microbiome Datasets Are Compositional: And This Is Not Optional.” Frontiers in Microbiology 8 (November):2224.

Hernández Medina, Ricardo, Svetlana Kutuzova, Knud Nor Nielsen, Joachim Johansen, Lars Hestbjerg Hansen, Mads Nielsen, and Simon Rasmussen. 2022. “Machine Learning and Deep Learning Applications in Microbiome Research.” ISME Communications 2 (1): 1–7.

Ibrahimi, Eliana, Marta B. Lopes, Xhilda Dhamo, Andrea Simeon, Rajesh Shigdel, Karel Hron, Blaž Stres, Domenica D’Elia, Magali Berland, and Laura Judith Marcos-Zambrano. 2023. “Overview of Data Preprocessing for Machine Learning Applications in Human Microbiome Research.” Frontiers in Microbiology 14 (October):1250909.

Kartal, Ece, Thomas S. B. Schmidt, Esther Molina-Montes, Sandra Rodríguez-Perales, Jakob Wirbel, Oleksandr M. Maistrenko, Wasiu A. Akanni, et al. 2022. “A Faecal Microbiota Signature with High Specificity for Pancreatic Cancer.” Gut 71 (7): 1359–72.

Kubinski, Ryszard, Jean-Yves Djamen-Kepaou, Timur Zhanabaev, Alex Hernandez-Garcia, Stefan Bauer, Falk Hildebrand, Tamas Korcsmaros, et al. 2022. “Benchmark of Data Processing Methods and Machine Learning Models for Gut Microbiome-Based Diagnosis of Inflammatory Bowel Disease.” Frontiers in Genetics 13 (February):784397.

Lagani, Vincenzo, Giorgos Athineou, Alessio Farcomeni, Michail Tsagris, and Ioannis Tsamardinos. 2017. “Feature Selection with the R Package MXM: Discovering Statistically Equivalent Feature Subsets.” Journal of Statistical Software 80 (September):1–25.

Lin, Xu, Hong-Mei Xiao, Hui-Min Liu, Wan-Qiang Lv, Jonathan Greenbaum, Rui Gong, Qiang Zhang, et al. 2023. “Gut Microbiota Impacts Bone via Bacteroides Vulgatus-Valeric Acid-Related Pathways.” Nature Communications 14 (1): 6853.

Liu, Yang, Guillaume Méric, Aki S. Havulinna, Shu Mei Teo, Fredrik Åberg, Matti Ruuskanen, Jon Sanders, et al. 2022. “Early Prediction of Incident Liver Disease Using Conventional Risk Factors and Gut-Microbiome-Augmented Gradient Boosting.” Cell Metabolism 34 (5): 719–30.e4.

Lopez-Siles, Mireia, Sylvia H. Duncan, L. Jesús Garcia-Gil, and Margarita Martinez-Medina. 2017. “Faecalibacterium Prausnitzii: From Microbiology to Diagnostics and Prognostics.” The ISME Journal 11 (4): 841–52.

Lundberg, Scott, and Su-In Lee. 2017. “A Unified Approach to Interpreting Model Predictions.” arXiv [cs.AI]. arXiv. http://arxiv.org/abs/1705.07874.

Marcos-Zambrano, Laura Judith, Kanita Karaduzovic-Hadziabdic, Tatjana Loncar Turukalo, Piotr Przymus, Vladimir Trajkovik, Oliver Aasmets, Magali Berland, et al. 2021. “Applications of Machine Learning in Human Microbiome Studies: A Review on Feature Selection, Biomarker Identification, Disease Prediction and Treatment.” Frontiers in Microbiology 12 (February):634511.

Martino, Cameron, James T. Morton, Clarisse A. Marotz, Luke R. Thompson, Anupriya Tripathi, Rob Knight, and Karsten Zengler. 2019. “A Novel Sparse Compositional Technique Reveals Microbial Perturbations.” mSystems 4 (1). 10.1128/mSystems.00016-19.

Nearing, Jacob T., Gavin M. Douglas, Molly G. Hayes, Jocelyn MacDonald, Dhwani K. Desai, Nicole Allward, Casey M. A. Jones, et al. 2022. “Microbiome Differential Abundance Methods Produce Different Results across 38 Datasets.” Nature Communications 13 (1): 342.

Pasolli, Edoardo, Lucas Schiffer, Paolo Manghi, Audrey Renson, Valerie Obenchain, Duy Tin Truong, Francesco Beghini, et al. 2017. “Accessible, Curated Metagenomic Data through ExperimentHub.” Nature Methods 14 (11): 1023–24.

Quinn, Thomas P., and Ionas Erb. 2020. “Interpretable Log Contrasts for the Classification of Health Biomarkers: A New Approach to Balance Selection.” mSystems 5 (2). 10.1128/mSystems.00230-19.

Ruuskanen, Matti O., Pande P. Erawijantari, Aki S. Havulinna, Yang Liu, Guillaume Méric, Jaakko Tuomilehto, Michael Inouye, et al. 2022. “Gut Microbiome Composition Is Predictive of Incident Type 2 Diabetes in a Population Cohort of 5,572 Finnish Adults.” Diabetes Care 45 (4): 811–18.

Salosensaari, Aaro, Ville Laitinen, Aki S. Havulinna, Guillaume Meric, Susan Cheng, Markus Perola, Liisa Valsta, et al. 2021. “Taxonomic Signatures of Cause-Specific Mortality Risk in Human Gut Microbiome.” Nature Communications 12 (1): 2671.

Silverman, Justin D., Kimberly Roche, Sayan Mukherjee, and Lawrence A. David. 2020. “Naught All Zeros in Sequence Count Data Are the Same.” Computational and Structural Biotechnology Journal 18 (September):2789–98.

Silverman, Justin D., Alex D. Washburne, Sayan Mukherjee, and Lawrence A. David. 2017. “A Phylogenetic Transform Enhances Analysis of Compositional Microbiota Data,” February. 10.7554/eLife.21887.

Singh, Sudha B., Amanda Carroll-Portillo, and Henry C. Lin. 2023. “Desulfovibrio in the Gut: The Enemy Within?” Microorganisms 11 (7). 10.3390/microorganisms11071772.

Venturelli, Ophelia S., Alex C. Carr, Garth Fisher, Ryan H. Hsu, Rebecca Lau, Benjamin P. Bowen, Susan Hromada, Trent Northen, and Adam P. Arkin. 2018. “Deciphering Microbial Interactions in Synthetic Human Gut Microbiome Communities.” Molecular Systems Biology 14 (6): e8157.

Wirbel, Jakob, Paul Theodor Pyl, Ece Kartal, Konrad Zych, Alireza Kashani, Alessio Milanese, Jonas S. Fleck, et al. 2019. “Meta-Analysis of Fecal Metagenomes Reveals Global Microbial Signatures That Are Specific for Colorectal Cancer.” Nature Medicine 25 (4): 679–89.

Wolpert, D. H., and W. G. Macready. 1997. “No Free Lunch Theorems for Optimization.” IEEE Transactions on Evolutionary Computation 1 (1): 67–82.

Zafar, Hassan, and Milton H. Saier Jr. 2021. “Gut Bacteroides Species in Health and Disease.” Gut Microbes 13 (1): 1–20.

Zeller, Georg, Julien Tap, Anita Y. Voigt, Shinichi Sunagawa, Jens Roat Kultima, Paul I. Costea, Aurélien Amiot, et al. 2014. “Potential of Fecal Microbiota for Early-Stage Detection of Colorectal Cancer.” Molecular Systems Biology 10 (11): 766.

Zeybel, Mujdat, Muhammad Arif, Xiangyu Li, Ozlem Altay, Hong Yang, Mengnan Shi, Murat Akyildiz, et al. 2022. “Multiomics Analysis Reveals the Impact of Microbiota on Host Metabolism in Hepatic Steatosis.” Advancement of Science 9 (11): e2104373.

Zhou, Yi-Hui, and Paul Gallins. 2019. “A Review and Tutorial of Machine Learning Methods for Microbiome Host Trait Prediction.” Frontiers in Genetics 10 (June):579.

Zou, Hui, and Trevor Hastie. 2005. “Regularization and Variable Selection Via the Elastic Net.” Journal of the Royal Statistical Society. Series B, Statistical Methodology 67 (2): 301–20.

